# Magnetic separation screens identify G alpha 15 as a central regulator of degranulation downstream of neutrophil chemoattractant receptors

**DOI:** 10.64898/2026.11.14.766678

**Authors:** Stefan M. Lundgren, Alexandra L. Calderon, Emel Akdoğan, Kimberly M. Bonger, Roarke A. Kamber, Michael C. Bassik, Sean R. Collins

## Abstract

Neutrophil recruitment and effector functions protect the body from infection. However, their excessive activity is central to inflammatory diseases, in which neutrophil granule proteins damage tissues and amplify inflammation. These proteins are released through degranulation, a regulated secretion process that can be triggered by the same chemoattractant molecules that drive neutrophil recruitment. Neutrophils detect attractants using G protein-coupled receptors (GPCRs) that signal through Gαi G-proteins. How the activity of these receptors is channeled to promote degranulation versus chemotaxis in the proper contexts remains unclear. Here, we identify positive and negative regulators of degranulation in HL-60 neutrophils using genome-wide CRISPR/Cas9 screens that exploit the transient surface exposure of granule membrane proteins. We identify Gα15 as a critical degranulation-specific effector of the attractant GPCRs FPR1 and LTB4R, and branched actin assembly as the dominant process limiting degranulation. Our results suggest that use of a second Gα subunit allows a switch to the degranulation output.

## Introduction

Regulated vesicle secretion is a fundamental eukaryotic cell behavior that enables cells to release molecules extracellularly and modify their membrane compositions. Its importance is highlighted in cellular processes including communication between neurons at synapses, insulin secretion by pancreatic beta cells, and targeted cell killing by T cells^1^. Multiple immune cell types, including neutrophils, use a regulated secretion process called degranulation to respond to stimuli by mobilizing receptors to their surface and releasing effector molecule^1–3^.

Neutrophils are innate immune cells that act as primary defenders against microbial infections and central mediators of inflammation^4,5^. Critical for these roles, they are highly adept at chemotaxis, the directed migration process through which they follow chemical signals to reach sites of infection, injury, and inflammation^5,6^. Upon reaching target locations, they respond to further stimulation with effector functions including degranulation, phagocytosis, and the formation of DNA-based extracellular traps. These effector functions kill and inactivate pathogens to control infections, but they also amplify inflammation and damage cells and tissues of the body^4,5,7^.

The proteins of neutrophil granules are central to their effector functions and to the related tissue damage. Neutrophils have multiple granule subtypes which have been classified according to the degree of cytotoxicity of their contents^3,8^. The less toxic granules subsets include additional receptors and other proteins that contribute to migration through tissues, and these granules are released more readily than the highly toxic primary granule subset. Primary granules include the myeloperoxidase enzyme, proteases, and antibacterial peptides. The release of these granules is the most highly regulated, requiring the strongest stimuli^3,8^. Elevated levels of primary granule proteins in blood and tissues are characteristic of severe reactions to COVID-19 and inflammation-associated diseases^9–12^.

Neutrophil chemotaxis is largely driven by a family of G protein-coupled receptors (GPCRs) that detect a range of attractants including pathogen-derived signals, chemokines, and signaling lipids^6^. The formyl peptide receptor FPR1 and the leukotriene B4 receptor LTB4R (also known as BLT1) are two receptors that play central roles in neutrophil recruitment and in excessive inflammatory reactions^13,14^. Notably, the same attractant receptors that direct chemotaxis, including FPR1 and LTB4R, can also trigger degranulation responses^3,15^. How these receptors can drive each behavioral output in the appropriate biological context remains incompletely understood.

Chemoattractant GPCRs primarily signal through the Gαi family of heterotrimeric G-proteins^6^. Gαi is thought to be essential for both the chemotaxis and degranulation responses, as both are blocked by pretreatment of cells with pertussis toxin, which inactivates Gαi^16^. Downstream of the receptors, the G-proteins activate signaling cascades that include the Rac and Cdc42 small GTPases, isoforms of phosphoinosite-3-kinase (PI3K), and phospholipase C beta (PLCβ). While Rac, Cdc42, and PI3K are central regulators of chemotaxis^17^, PLCβ may be more important for degranulation (Fig. 1a). Across multiple immune cell types and stimuli, phospholipase C activities lead to protein kinase C (PKC) and calcium signals that are critical for degranulation^2,18–20^. Downstream of these signals, the GTPase Rab27a coordinates granule secretion, working with the calcium-dependent tethering protein Munc13-4^21,22^. Even so, much remains unknown about the mechanisms connecting the upstream signals to granule mobilization and secretion, the mechanisms restricting the release of primary granules, and the mechanisms controlling the channeling of GPCR activity towards degranulation versus chemotaxis responses.

**Fig. 1:**
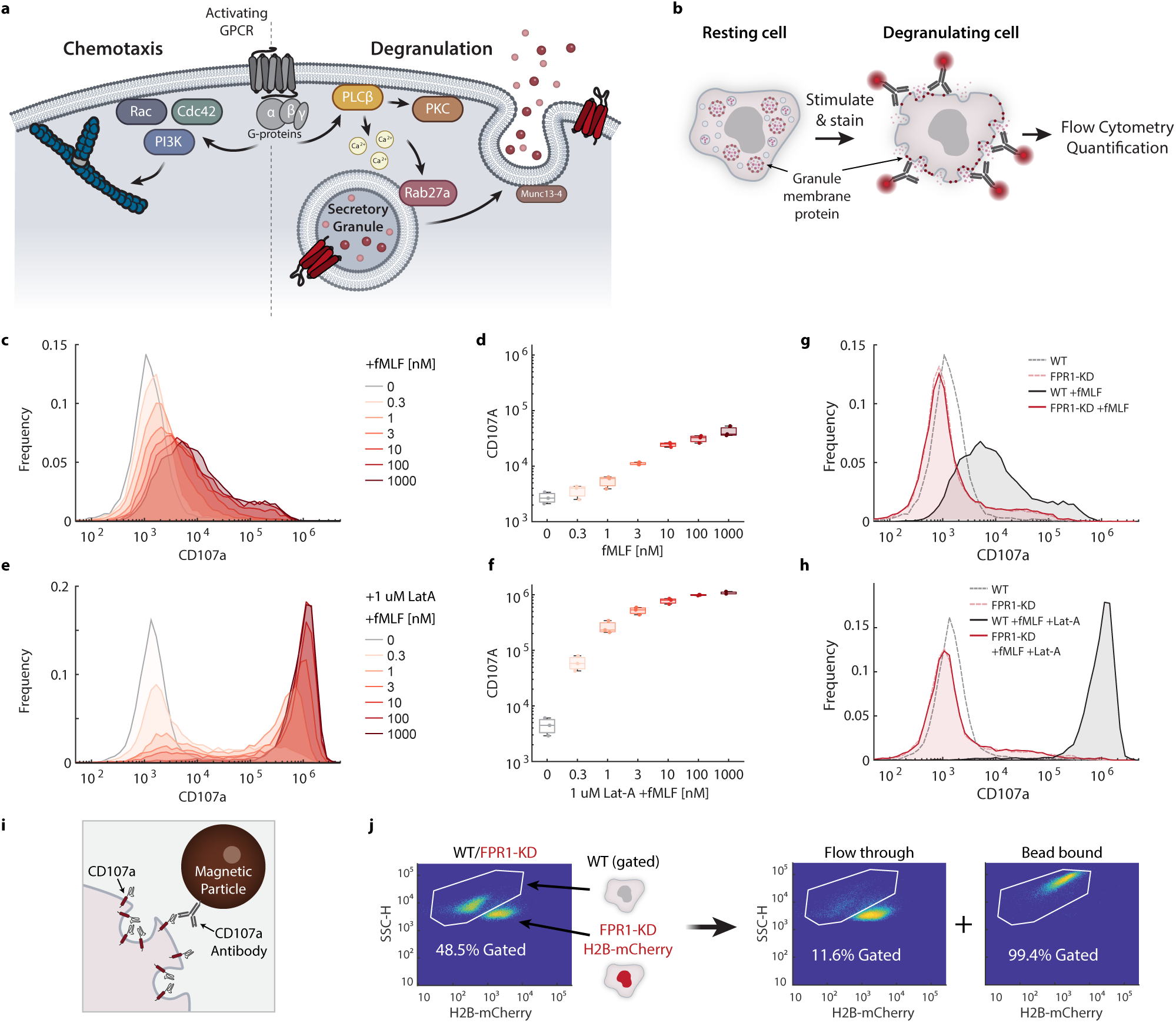
FPR1 induces degranulation and dose-dependent surface expression of granule membrane proteins. **a,** Schematic describing simplified versions of neutrophil receptor signaling cascades leading to dual chemotaxis and degranulation outputs. **b,** Schematic illustrating quantification of degranulation by flow cytometry. Granule membrane proteins are not present on the surface of a resting cell, but they become exposed on the surface due to membrane fusion during granule exocytosis. Fluorophore-conjugated antibodies can be used to detect changes in the surface exposure of these proteins as a measure of degranulation. **c,** Representative histograms showing cell surface levels of the granule membrane protein CD107a following treatment with fMLF in dHL-60 cells. **d,** Boxplots quantifying mean CD107a surface levels as an fMLF dose response. Dots indicate independent biological replicates. **e,** Representative histograms of CD107a surface levels in wild-type dHL-60 cells before and after stimulation with the indicated concentrations of fMLF after pretreatment with Latrunculin-A (LatA) for 30 minutes and **f,** corresponding boxplots showing biological replicate data. **g-h,** CD107a surface levels, comparing wild type and CRISPRi FPR1 knockdown (FPR1-KD) dHL-60 cells in response to fMLF or LatA and fMLF treatment. **i,** Schematic illustrating the use of antibody-coated magnetic particles to capture cells based on surface expression of the granule membrane protein CD107a. **j,** Magnetic separation of wild-type from FPR1-KD cells based on CD107a cell surface expression after treatment with LatA and 1 uM fMLF. Wild-type cells expressing no fluorescent proteins were mixed in a 1 to 1 ratio with FPR1-KD cells expressing Histone H2B-mCherry. Density-colored scatter plots from cytometry measurements show cell populations before and after separation. The gate shown was used to measure the percent wild-type cells in each population. For all figures, boxes in boxplots indicate the 25th and 75th percentiles, with the center bar indicating the median and whiskers indicating the range of the data aside from automatically determined outliers.

Here, we identify positive and negative regulators of GPCR-driven degranulation using parallel genome-wide CRISPR/Cas9 screens in neutrophil-differentiated HL-60 (dHL-60) cells^23,24^. To accomplish this, we develop and apply a screening strategy using immuno-magnetic cell separation based on the exposure of granule membrane proteins on the cell surface. Our screens uncover many known and previously unknown components regulating degranulation. We identify branched actin assembly, mediated by the WAVE and Arp2/3 complexes, as a dominant pathway limiting degranulation. Furthermore, we identify Gα15 as a degranulation-specific effector that is critical for the release of primary granules and CD107a-marked organelles downstream of both FPR1 and LTB4R. Our results suggest that engagement of Gα15 provides a key signal that shifts chemoattractant responses to the degranulation output behavior.

## Results

### Formyl peptide induced degranulation is dose- and FPR1-dependent

We sought to develop a screening approach to systematically identify genes regulating degranulation, and we reasoned that cell surface exposure of granule membrane proteins could be leveraged for selection. These proteins are normally hidden from the cell surface, but they become exposed when the granule fuses with the plasma membrane during degranulation. Their abundance on the cell surface can be measured by flow cytometry, and exocytosis of different granule subtypes can be monitored using corresponding granule protein markers^23,25^.

To understand how chemoattractant receptors control degranulation, we focused on FPR1 as a model attractant GPCR, and dHL-60 cells as a genetically tractable model for human neutrophils. We used CD107a (also known as LAMP1) as a degranulation marker with very low basal surface expression and a large increase after stimulation^26^ – a key feature for a successful screen. CD107a is used as a marker for degranulation across multiple immune cell types, including T cells and NK cells^23,27,28^. In neutrophils and dHL-60 cells, it localizes to mobilizable organelles including granules and multivesicular bodies^23,26,29^. To measure degranulation, we stained cells before and after stimulation with anti-CD107a antibodies conjugated to a fluorophore (Fig. 1b).

Consistent with prior studies, we found that stimulating dHL-60 cells with the FPR1 ligand N-formyl-methionine-leucyl-phenylalanine (fMLF) results in a dose-dependent degranulation response (Fig. 1c-d). However, the response is greatly amplified when the cells are pretreated with Latrunculin A (LatA) (Fig. 1e-f). LatA causes actin disassembly in cells by sequestering actin monomers, and it is thought to facilitate degranulation by removing a barrier of cortical actin^25,30^. Furthermore, particularly in the presence of LatA, the response to fMLF was bimodal at the population level. Individual cells either induced strong degranulation or did not degranulate, but the fraction of degranulating cells increased with increasing stimulus concentration (Fig. 1c-f).

Next, we verified that the degranulation responses were dependent on the receptor FPR1. We generated a cell line with stable knockdown of FPR1 through CRISPR interference (CRIPSRi). We observed a near complete reduction in degranulation in FPR1 knockdown cells, both in the presence and absence of LatA (Fig. 1g, h).

### Magnetic separation of degranulating cells using a CD107a molecular handle

While fluorescence activated cell sorting can be used for genome-wide CRISPR screens, we reasoned that the large change in cell surface CD107a could allow for rapid magnetic sorting with higher throughput. With this goal in mind, we tested whether CD107a could be used as a molecular handle to physically separate cells that had degranulated from those that had not (Fig. 1i). As a proof-of-principle experiment, we mixed wild-type dHL-60 cells with FPR1 knockdown (FPR1-KD) cells at an approximately 1 to 1 ratio. We included a Histone H2B-mCherry marker in the FPR1-KD cells to allow us to track the two populations. We stimulated the mixed population of cells with fMLF and LatA, and then used magnetic beads coated in anti-CD107a antibodies to separate cells. Quantifying by flow cytometry, we found that more than 99% of the cells in the bead-bound fraction were wild-type cells (Fig. 1j). In contrast, less than 12% of cells in the flow through fraction were wild-type cells. Our results indicate that immuno-magnetic separation using a CD107a molecular handle can effectively separate cells based on a degranulation phenotype.

### Parallel genome-wide screens for degranulation and differentiation

Using our separation strategy, we implemented two genome-wide CRISPR/Cas9 screens for fMLF-stimulated degranulation (Fig. 2a). One screen included pretreatment with LatA, which leads to high rates of degranulation (Fig. 1e-f). We reasoned that this screen would be sensitized to detect genes required for degranulation, although it could miss genes related to actin remodeling. The second screen included only fMLF stimulation, which results in lower rates of degranulation (Fig. 1c-d), making it better suited to detect the components limiting degranulation when the actin cytoskeleton is intact, and the components needed to modify the cytoskeleton to promote degranulation. Since neutrophil differentiation is required for degranulation, we also completed a third screen for differentiation, following a strategy used previously with CRISPR interference^31^. We reasoned that this screen could be used to distinguish genes that directly impact the degranulation process from those that affect degranulation as a secondary consequence of an effect on differentiation.

**Fig. 2:**
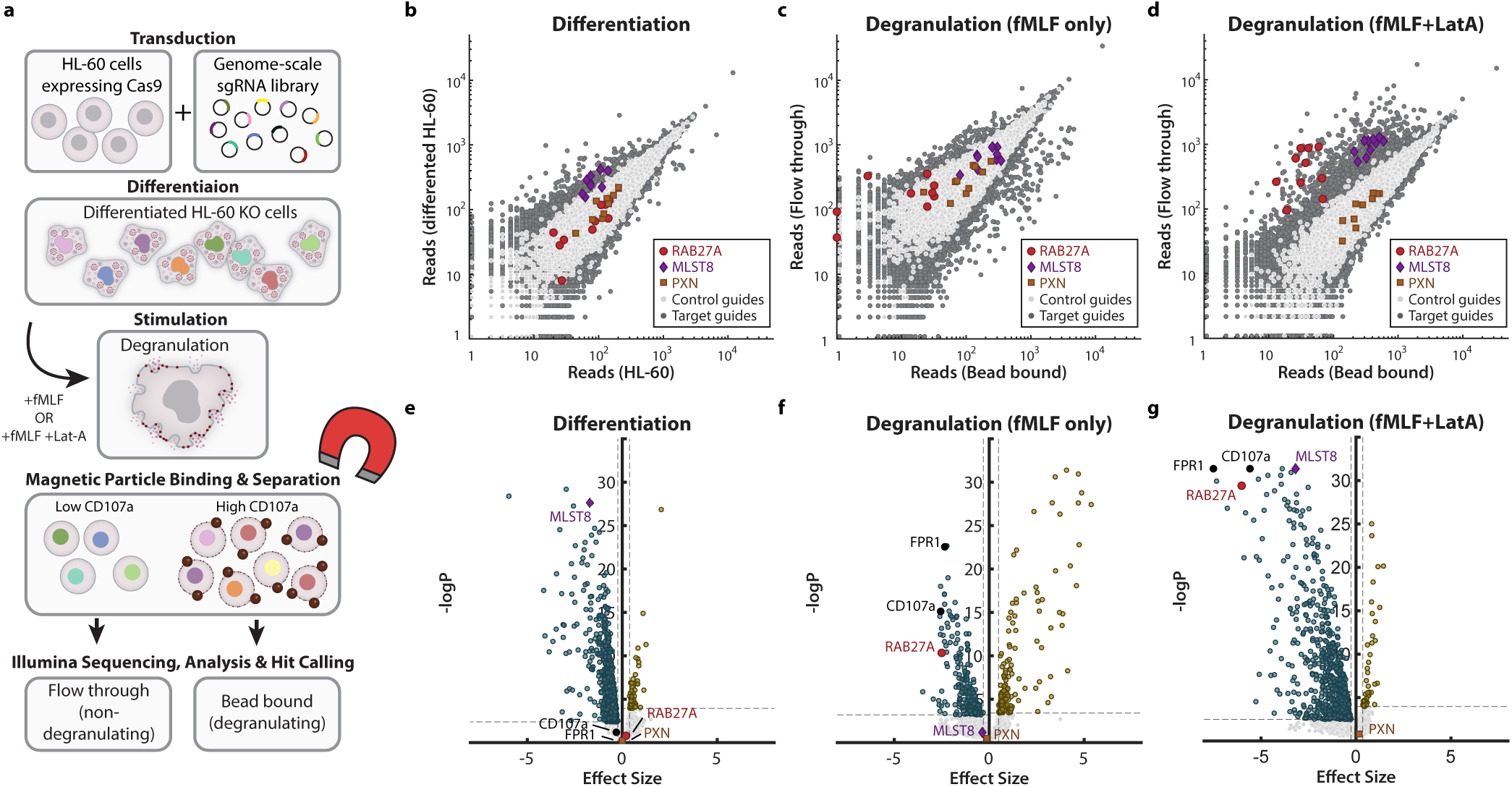
Pooled genome-wide CRISPR/Cas9 knockout screens for differentiation and degranulation. **a,** Schematic of the workflow for pooled CRISPR screens to identify regulators of degranulation in neutrophil-like cells. The genome-scale library contains 10 sgRNA sequences per gene in the human genome and ∼12,000 negative control sgRNA sequences. Undifferentiated and differentiated HL-60 cells were each collected for a differentiation screen. For the degranulation screens, dHL-60 cells were stimulated with 1 uM fMLF or incubated for 30 min with LatA and then stimulated with 1 uM fMLF prior to magnetic cell separation. sgRNAs were sequenced for bead bound and flow through cell populations. Sequencing results were analyzed to compute log-likelihood scores, p-values, effect sizes, and a signed logP score for each gene (see Methods for details). **b-d,** Scatter plots showing sgRNA sequence read count comparisons for three separate screens to identify genes involved in differentiation and degranulation. Each of the 10 sgRNAs are highlighted for the representative genes RAB27A, MLST8, and PXN. Negative control sgRNAs are shown in light gray. Other sgRNAs are shown in dark gray. **e-g,** Volcano plots showing calculated effect size estimates against negative log_10_ p-values for each gene. Genes below a 5% false discovery rate threshold are shown in dark cyan (calculated for negative score) or gold (calculated for positive scores).

To implement our screens, we used HL-60 cells stably expressing Cas9 and transduced with a genome-wide CRISPR knockout library containing 10 unique sgRNAs for each gene, as well as more than 12,000 non-targeting and safe-targeting control sgRNAs^32^. The cells also express an exogenous copy of FPR1 to reduce heterogeneity in receptor expression. We previously used this cell line and library for screens to identify regulators of FPR1 trafficking and surface expression^33^, and we verified a high level of Cas9 activity using a self-targeting GFP construct (Extended Data Fig. 1a). We differentiated the pooled knockout cell library into a neutrophil-like state^24^ and split the cells into three pools of ∼4.0×10^8^ cells. For our differentiation screen, we sequenced one pool and compared it to the undifferentiated HL-60 knockout library. We used the other pools for the two degranulation screens. After stimulation, we magnetically separated cells into bead-bound (degranulating) and flow through (non-degranulating) fractions and used sequencing to compare the abundance of each guide between the populations (Fig. 2a).

As an initial test of the effectiveness of our screens, we analyzed the sequence read data for three well-characterized genes: RAB27A, which is a critical and specific regulator of degranulation^21,22^, MLST8, which is needed for cell-cycle arrest during differentiation^31^, and Paxillin (PXN), which was not expected to be needed for differentiation or fMLF-induced degranulation. The results aligned well with these expectations. The guides targeting RAB27A were enriched in the non-degranulating populations in our degranulation screens but behaved similarly to the control guides in the differentiation screen (Fig. 2b-d). The guides targeting MLST8 were outliers in the differentiation screen (Fig. 2b), and consistent with our rationale for that screen, these guides were also outliers for the fMLF+LatA screen, likely as a secondary consequence of the differentiation defect (Fig. 2d). Finally, the guides targeting PXN behaved similarly to control guides in all three screens (Fig. 2b-d).

To analyze the screen data formally, we used the CRISPR Log Odds (CLO) approach we previously developed which provides guide-level (Supplementary Data 1) and gene-level scores (Supplementary Data 2), including log-likelihood scores based on a negative binomial model, empirical p-values based on rank-order relative to control guides, and Bayesian effect size estimates^33^. From the p-values, we computed false discovery rate (FDR) estimates, as well as a “logP score” which is equal to the negative log10 p-value with a sign (positive or negative) indicating the direction of the effect. For the degranulation screens, negative scores correspond to decreased degranulation. For the differentiation screen, a negative score corresponds to increased abundance in the differentiated cell population, which may represent a defect in differentiation-associated cell cycle arrest. Finally, we defined hit genes using an FDR threshold of 5% (Fig. 2e-g).

As an additional test of the specificity of our results, we compared the screen scores for each gene to the corresponding mRNA gene expression levels in dHL-60 cells^24^. We reasoned that genes that are not expressed should not affect degranulation or differentiation and would effectively serve as additional negative controls. Consistent with this expectation, we found that genes with low or undetected mRNA levels had scores tightly distributed near zero (Extended Data Fig. 1b-f). Additionally, we observed a trend in which genes with higher expression levels were much more frequently identified as hits (Extended Data Fig. 1c).

### Genome-wide screens reveal novel regulators of neutrophil degranulation

We identified many hits for each degranulation screen, and we wanted to focus our attention on the pathways directly controlling neutrophil degranulation. Given that differentiation is important for forming granules and setting proper expression levels of proteins involved in signaling, we reasoned that genes required for differentiation would often have secondary effects on degranulation. Indeed, many genes showed correlated effects on differentiation and degranulation (Supplementary Data 2). To identify the genes most likely to be direct regulators of degranulation, we created a narrower list of degranulation hits, excluding genes that passed a 5% FDR cutoff for differentiation. We included genes that were hits in either degranulation screen, resulting in a list of 846 degranulation-specific hits (Supplementary Data 3).

To get more insight into the pathways controlling degranulation, we manually annotated the hits into functional categories based on prior literature (Fig. 3, Extended Data Fig. 2, Supplementary Data 3). In some cases, we included genes that were moderate differentiation hits to get a more complete perspective of the hits in each category. We found that our screen hits effectively identified known regulators of neutrophil degranulation, including the receptor (*FPR1*), Gαi heterotrimeric G-protein subunits (*GNAI2*, *GNB1*, *GNB2*, and *GNG5*), the major PLC isoform downstream of GPCRs in these cells (*PLCB2*), the IP3 receptor (*ITPR2*), multiple isoforms of PKC (*PRKCA*, *PRKCB*, and *PRKCD*), store-operated calcium signaling components (*STIM1* and *ORAI1*), RAC2 and its regulator DOCK2, the actin disassembly factors Cofilin and Coronin (*CFL1* and *CORO1A*), and direct regulators of secretion including the Rab 27a GTPase (*RAB27A*), its effector protein Munc13-4 (*UNC13D*), the SNARE protein SNAP23, and the fusion regulator STXBP2 (also known as Munc18-2)^18,34,35^.

**Fig. 3:**
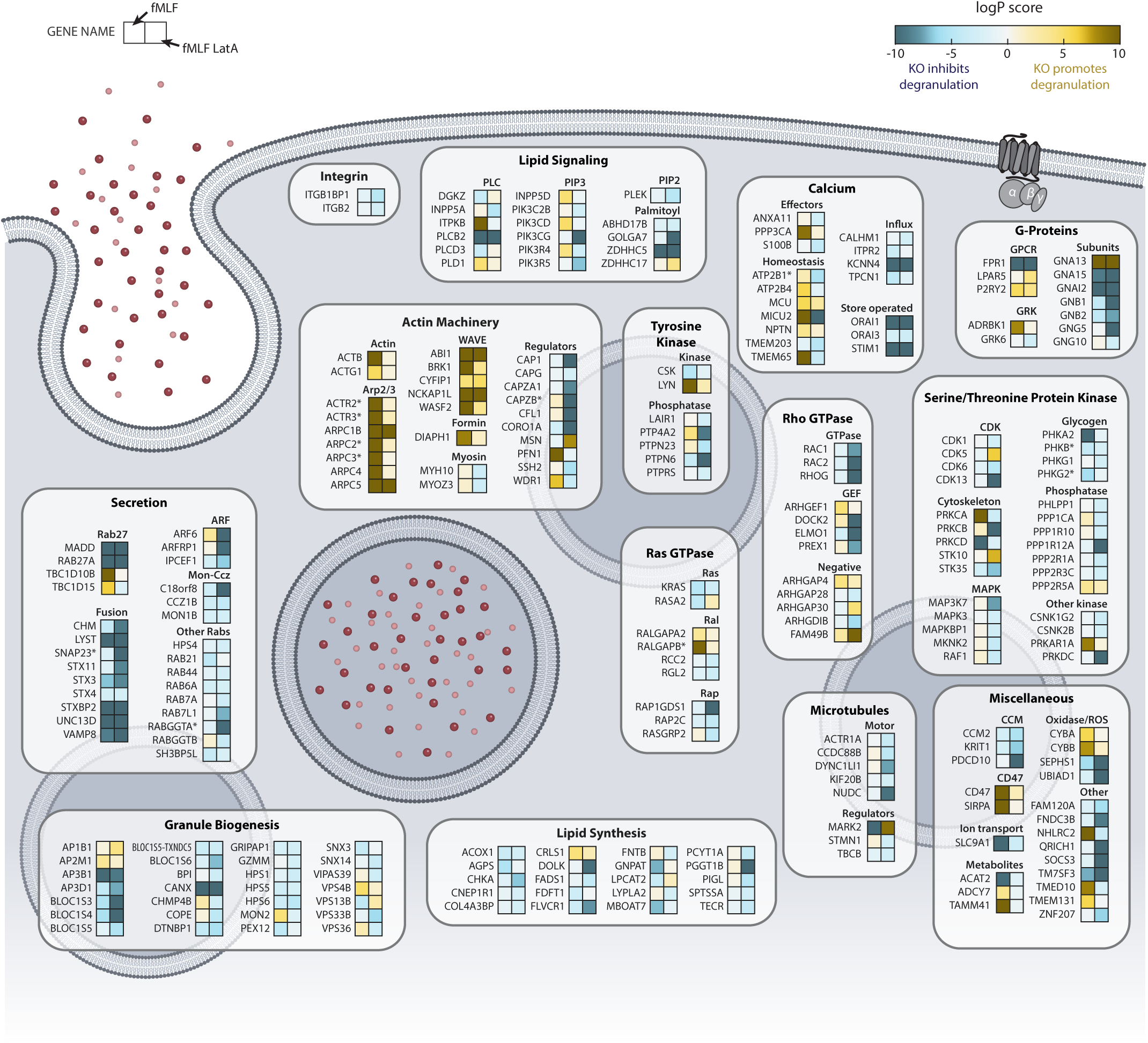
Overview of hits from genome-wide degranulation screens. The schematic shows logP score heatmaps for selected hit genes, organized by functional categories. All genes shown passed a 5% FDR threshold for at least one degranulation screen. Genes that were also hits in the differentiation screen were excluded, unless marked with (*). These select differentiation and degranulation hits were included here to highlight data set trends. For each gene, the score for the fMLF only screen is shown on the left, and the score for the screen with fMLF and LatA is shown on the right. Functional categories were assigned manually based on prior literature.

We also identified multiple components that may fill gaps in our understanding of neutrophil degranulation. While Rab27a acts as a master regulator of degranulation, its direct regulators in neutrophils had not been characterized^18,22^. We identified MADD and TBC1D10B, which have been implicated as a GEF and GAP, respectively, for Rab27a in other cell types^28,36,37^ (Fig. 3). Consistent with these proposed roles, MADD is required for degranulation and TBC1D10B limits degranulation. We further identified components of the secretory machinery and components involved in G-protein, lipid, calcium, Rho GTPase, Tyrosine kinase, and Serine/Threonine kinase signaling pathways (Fig. 3). We identified specific components of both the actin and microtubule cytoskeletons, and components involved in granule biogenesis (Fig. 3).

### Exocytosis of primary granules and CD107a-marked organelles share mechanistic control

Neutrophil granules consist of multiple subtypes that vary in their cargo, and granule exocytosis is ordered, with granules containing more cytotoxic content requiring higher levels of stimulus for release^38^. Secretory vesicles, which largely mobilize additional receptors to the cell surface, are the most readily exocytosed. These are followed by secondary, gelatinase-containing, and ficolin-rich granules^26,38^. Primary granules are the most toxic and the last to be released. Additionally, neutrophils can exocytose multivesicular bodies, a process linked to exosome release and the production of LTB4 gradients that amplify neutrophil recruitment, in a process similar to degranulation^39,40^.

Our degranulation screen results provided an opportunity to assess the overlap between the mechanisms controlling release of different granule types. Granule subtypes and their secretion can be identified by membrane protein markers, with CD63 marking primary granules, CD11b marking intermediate granule types, and CD35 marking secretory vesicles (Extended Data Fig. 3)^23,41^. Our screen marker, CD107a has been reported to localize to mobilizable organelles that are distinct from primary granules, including multivesicular bodies^29,42^. However, in a proteomic study it was found on primary granules, multivesicular bodies, and also other granule subtypes to a lesser extent^26^. As further validation of our screen results, and to determine the degree to which mechanisms and components are shared across different secretory organelles, we created individual CRISPRi knockdown cell lines for a set of 23 degranulation-specific hit genes (Fig. 4a, b). For each gene, we made two independent cell lines with distinct sgRNA guide sequences (Supplementary Table 1). Then, for each knockdown cell line, we measured the stimulated increase in surface expression of four marker proteins: CD107a, CD63, CD11b, and CD35, as well as the surface levels of FPR1 (Fig. 4c, Extended Data Fig. 4-5).

**Fig. 4:**
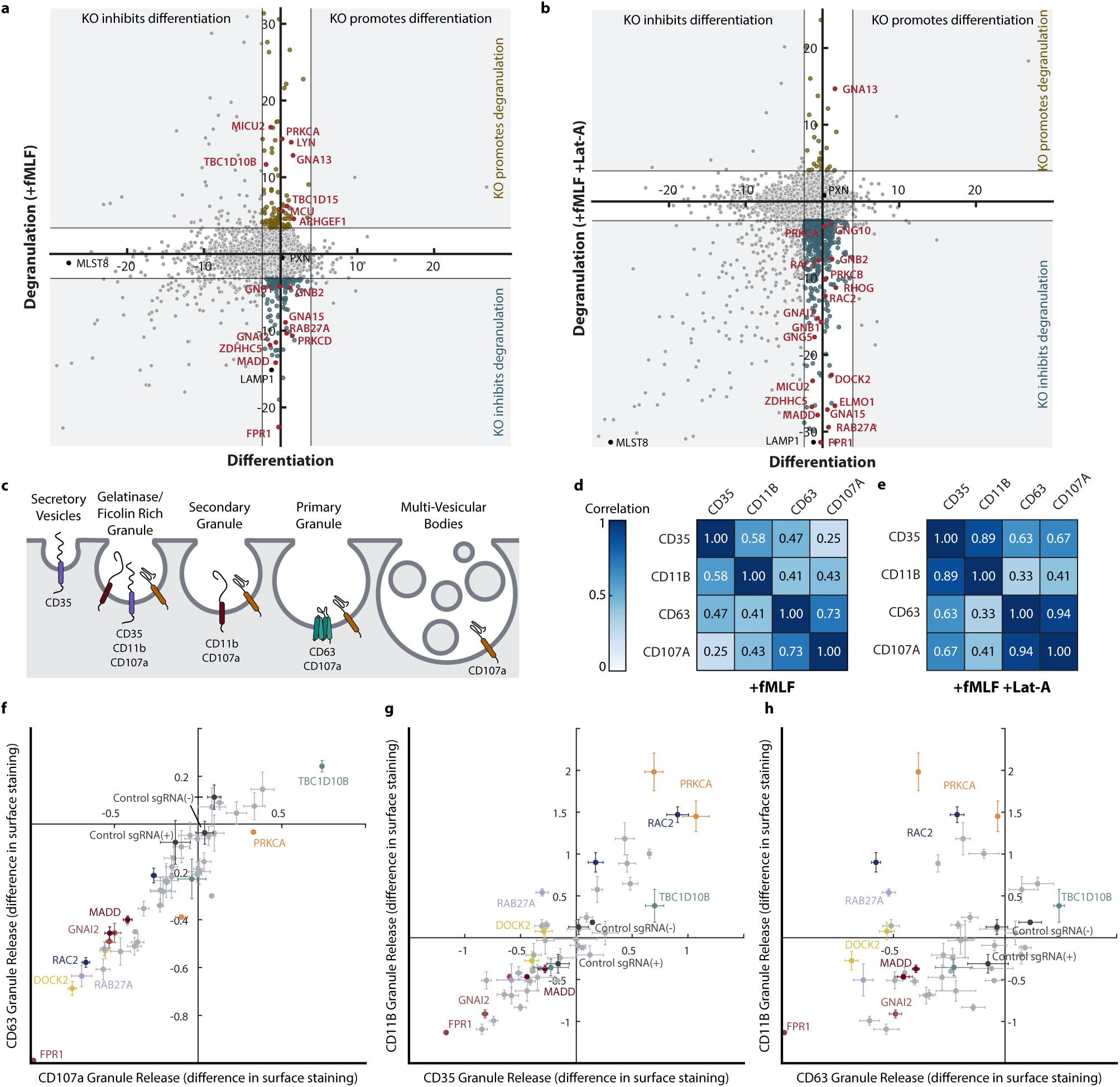
Degranulation hits differentially affect exocytosis of different granule subtypes. **a**-**b,** Scatter plots displaying logP scores for each gene, comparing the differentiation screen on the x-axis to the indicated degranulation screen on the y-axis. Gray lines near the axes indicate the 5% FDR significance threshold for the positive and negative side of each screen. Gene knockouts that affected degranulation, but yielded no significant difference in differentiation, are highlighted in dark cyan (decreased degranulation) or gold (increased degranulation). LAMP1 (CD107a), the granule membrane marker used for physical separation of cells, is highlighted in black. A select set of hits shown in red were chosen for follow up experiments. For each of these genes, two separate CRISPRi knockdown cell lines were created. **c,** Schematic showing an approximate distribution of commonly used granule markers across granule sub-types. **d**-**e,** Matrix of Pearson’s correlation coefficients comparing the effects of the knockdown of the selected genes on exocytosis of different granule subtypes. The effects were measured as the normalized change in cell surface expression of the indicated granule membrane proteins (CD35, CD11b, CD63, and CD107a) after stimulation. Correlation matrices are shown for responses to stimulation with 100 nM fMLF (**d**) or 10 nM fMLF and 1 uM LatA (**e**). **f-h**, Scatter plots showing the effects of gene knockdowns on the degranulation of the indicated granule sub-types, corresponding to data shown in **e**. Data are normalized to the mean of three control samples. Data points and error bars represent the mean and ±s.e.m. for n=3 independent biological replicates.

We found that the effects of gene perturbations on the exocytosis of CD107a and CD63 were tightly correlated (Fig. 4d-f). Indeed, the correlation coefficient for these two markers for the fMLF+LatA stimulus was 0.94 (Fig. 4e-f). Consistent with their putative roles as positive and negative regulators of Rab27a, we observed that MADD was required for degranulation of CD63 granules, while TBC1D10B limited their release (Fig. 4f). Our results suggest that the mechanisms controlling the release of primary granules and CD107a-marked mobilizable organelles are largely overlapping.

We also observed tight correlations for the effects of gene perturbations on the secretion of CD11b- and CD35-marked granules (Fig 4d, e, g). However, the effects on these two markers were less correlated with effects on CD107a- or CD63-marked organelles (Fig. 4d, e, h). Consistent with reports that Rab27a regulates primary granules, but not other granule subsets, we observed only mild phenotypes for RAB27A, MADD, and TBC1D10B for secretion of CD11b and CD35. More generally, phenotypes for multiple genes differed between the CD63 and CD11b granule subtypes (Fig. 4h). The different components and mechanisms involved for the granule classes may contribute to their different sensitivity for release.

### Branched actin assembly is a dominant process restraining degranulation

Our screen without LatA provided an excellent opportunity to identify components and pathways limiting degranulation in response to GPCR inputs. Indeed, we identified many such components from multiple pathways (Fig. 3). These components included inositol-trisphosphate 3-kinase B (*ITPKB*) which phosphorylates IP3 to terminate activation of IP3 receptors, two components of the NADPH oxidase (*CYBA* and *CYBB*) which can oppose calcium influxes^43^, and the putative Rab27a GAP TBC1D10B. However, many of the strongest hits for genes limiting degranulation were regulators of the actin cytoskeleton (Fig. 3, Supplementary Data 2).

Cortical actin under the plasma membrane is thought to act as a barrier that limits degranulation in neutrophils and other immune cells^25,44^. However, the cortical actin network involves both linear and branched actin, which is assembled through the combined activity of formin- and Arp2/3-mediated nucleation, as well as filament crosslinking, contractile pulling by myosin II, and plasma membrane attachment through ERM proteins. Among these protein classes, we found that the formin mDia1 (*DIAPH1*), the ERM protein moesin (*MSN*), and the ERM-protein activator LOK (*STK10*) all contribute to limiting degranulation (Fig. 3). However, of the 11 strongest hits in our screen for genes limiting degranulation, 6 were components of the Arp2/3 complex (*ACTR2*, *ACTR3*, *ARPC1B*, *ARPC2*, *ARPC3*, and *ARPC5*) and 4 were components of the WAVE2 complex (*WASF2*, *NCKAP1L*, *BRK1*, and *ABI1*). We did not find WASP or other Arp2/3 activators among the screen hits. These results suggest that branched actin assembly driven by the WAVE complex is the dominant pathway limiting degranulation.

To validate our findings and confirm the importance of branched actin in restraining degranulation, we tested whether chemical inhibitors of Arp2/3 (CK666 and CK869)^45,46^, formins (SMIFH2)^47^, and the SLK/LOK kinases that activate ERM proteins (SLK/STK10-in-1, also known as Cpd31)^48^ facilitate degranulation. We pretreated cells with chemical inhibitors, stimulated them with fMLF, and measured degranulation responses using both the CD107a and CD63 markers. Among these inhibitors, only the combined Arp2/3 inhibitors caused increased degranulation (Fig. 5a-d). Even so, the effect of this treatment was noticeably weaker than that of LatA. In contrast, SMIFH2 caused a slight, but significant decrease in degranulation.

**Fig. 5:**
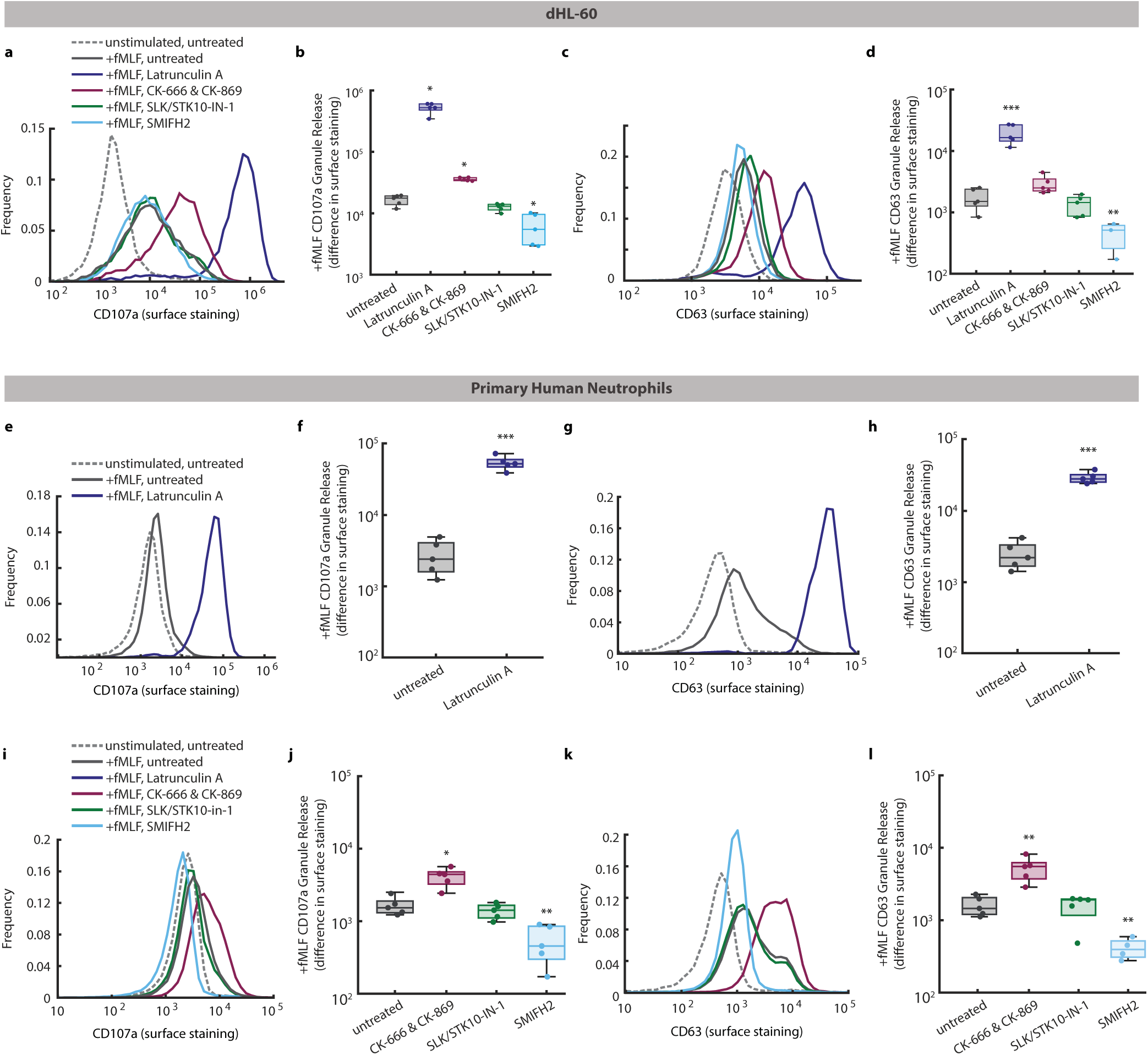
Branched actin restrains degranulation. **a-d**, Comparison of the degranulation response of dHL-60 cells stimulated with 100 nM fMLF in the presence or absence of the actin depolymerizing agent LatA, the Arp2/3 inhibitors CK-666 and CK-869, the SLK/LOK inhibitor SLK/STK10-IN-1, or the formin inhibitor SMIFH2. Representative histograms show the **(a)** CD107a and **(c)** CD63 surface expression for either untreated cells or cells pretreated with the indicated actin inhibitors for 30 minutes and stimulated with fMLF. Corresponding boxplots show the mean increase in **(b)** CD107a and **(d)** CD63 surface levels after stimulation with fMLF. **e-h,** Comparison of the degranulation response of primary human neutrophils stimulated with 100 nM fMLF with or without pretreatment with LatA for 30 minutes. Representative histograms show the cell surface levels of **(e)** CD107a and **(g)** CD63. Corresponding boxplots show the mean increase in **(f)** CD107a and **(h)** CD63 surface levels after stimulation with fMLF. **i-l,** Representative histograms and boxplots for primary human neutrophils stimulated with 100 nM fMLF after pretreatment with the indicated actin inhibitors for 30 minutes. In all boxplots, dots indicate independent biological replicates, with n=5 independent biological replicates for all samples except for dHL-60 cells treated with SMIFH2, for which n=3. P-values were calculated for each condition using log-transformed values, comparing against the no inhibitor control condition using ANOVA and Tukey’s multi-comparison test (panels d,f), T-test (panels h,i), or Mann-Whitney with the Bonferonni correction (panel b). For all figures, * indicates a multiple comparison corrected p-value of < .05, ** indicates < .01, and *** indicates < .001.

We further confirmed the effects of these inhibitors in primary human neutrophils. While gene knockdowns are not possible in these cells due to their short lifespan, we could perform experiments using the inhibitors. Our results in primary human neutrophils paralleled our results with dHL-60 cells. Stimulation of primary neutrophils with fMLF after LatA treatment induced a strong degranulation response, detectable with both the CD107a (Fig. 5e-f) and CD63 (Fig. 5g-h) markers. Of the other inhibitors, only the combined Arp2/3 inhibitors increased degranulation (Fig. 5i-l). This effect was significant for both CD107a and CD63. Also consistent with our results in dHL-60 cells, SMIFH2 caused decreased degranulation (Fig. 5i-l). Together, our results with actin network inhibitors support our screen results, indicating that branched actin assembly is a dominant process limiting degranulation.

### GNA15 is a component that distinguishes control of chemotaxis and degranulation downstream of FPR1

A primary goal of our study was to identify components that may control the channeling of receptor inputs to the degranulation response, rather than chemotaxis. Towards this aim, we used our previously developed 96-well format chemotaxis assay^49^, based on automated photo-uncaging of fMLF, to measure migration and chemotaxis phenotypes for the same focused set of CRISPRi knockdowns that we used to analyze the release of different mobilizable organelles (Fig. 6a).

**Fig. 6:**
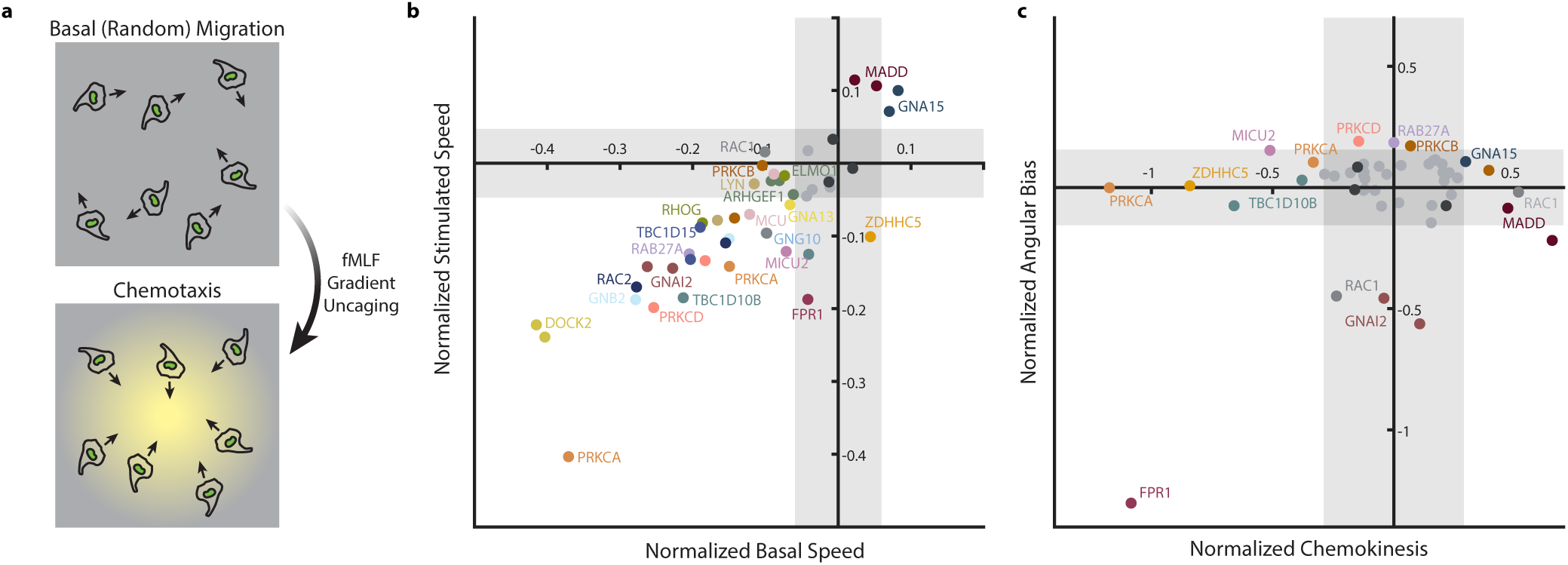
Many degranulation hit genes, but not GNA15, are needed for directed cell migration. **a,** Schematic of chemotaxis assay based on attractant uncaging. Live-cell imaging was used to track the migration of dHL-60 cells before and after generation of an attractant gradient by photo-uncaging of fMLF in 96-well imaging plates. Migration speed and directionality parameters were computed from cell tracks. **b,** Scatter plot comparing mean phenotype scores for basal and post-stimulus cell speed for each CRISPRi knockdown cell line (2 independent guide sequences per gene). Values were normalized to the mean of three control cell lines. Controls, highlighted with black dots, were two technical replicates of wild type cells and one safe targeting sgRNA. Each point represents the mean of n=3 independent biological replicates. The gray-shaded regions indicate 98% confidence intervals based on fitting of the empirical error distribution. **c,** Scatter plot of mean phenotype scores for chemokinesis and angular accuracy.

We found that knocking down many of these genes resulted in defects in cell migration, with knockdown cells migrating slower than control cells both before and after generation of a chemoattractant gradient (Fig. 6b, Supplementary Data 4). Consistent with previous findings, RAC2 and DOCK2, an activator of Rac GTPases, had particularly strong migration defects (Fig. 6b)^49^. These results suggest that DOCK2 acts as a central activator of Rac GTPases for both the migration and degranulation behaviors.

Knockdown of *GNAI2*, which encodes the major Gαi isoform in leukocytes, caused a strong defect in the directional accuracy of chemotaxis (Fig. 6c). This result is consistent with the central role of Gαi in chemotaxis and prior studies demonstrating directional migration defects when *GNAI2* is knocked down or deleted^31,49–51^.

In contrast, we found other degranulation hit genes that had little effect on migration or chemotaxis (Fig. 6b-c). Notably, *GNA15* (also previously known as *GNA16*) was one of the strongest hits in terms of genes required for degranulation in our screens (Fig. 4a,b), and we found that knockdown of *GNA15* had no deleterious effect on chemotaxis. Indeed, it resulted in slightly elevated migration speed with no defect in directionality (Fig. 6b-c). These results are consistent with chemotaxis results from our prior siRNA screen and from a study in mouse macrophages^49,50^, and they suggest that Gα15 may be a degranulation-specific effector acting immediately downstream of FPR1.

### G*α*15 controls degranulation downstream of GPCRs

We then asked whether Gα15 is a mediator of degranulation specifically for formyl peptides, or whether it is required for degranulation in response to other inputs as well. We considered LTB4 as a second model GPCR ligand and opsonized zymosan as an input that acts through non-GPCR receptors. Opsonized zymosan can trigger phagocytosis and degranulation in neutrophils, acting in large part through Complement Receptor 3 (CR3, also known as Mac-1 or CD11b/CD18 integrin)^52,53^.

We first verified that each input is capable of triggering degranulation. In contrast to formyl peptides, we found that treatment of dHL-60 cells with LTB4 alone resulted in negligible amounts of degranulation (Fig. 7a). This result is consistent with the more transient signaling output of LTB4R, relative to that of FPR1^54^. However, when LTB4 was added to cells pretreated with LatA, it induced robust degranulation (Fig. 7b). Stimulation with human serum-opsonized zymosan alone was sufficient to induce degranulation at levels close to that induced by fMLF (Fig. 7a).

**Fig. 7:**
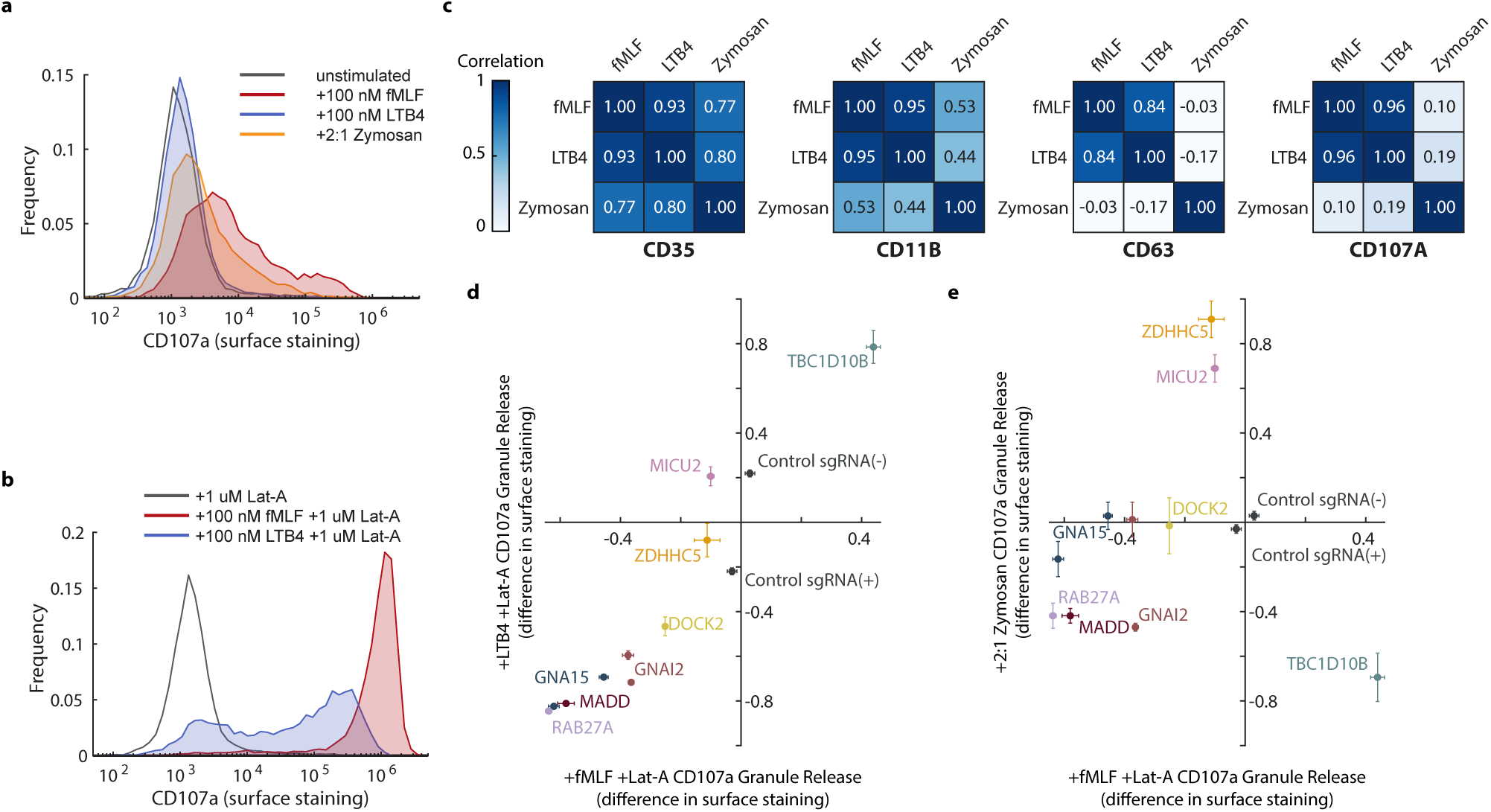
GPCRs share a specific degranulation pathway. **a,** Histograms comparing dHL-60 degranulation responses to different stimuli as measured by CD107a cell surface expression. For the Zymosan stimulation condition, human serum opsonized Zymosan A S. *cerevisiae* particles were added at a 2 particle to 1 cell ratio. **b,** Histograms comparing the degranulation response to fMLF or LTB4 when pretreated with 1 uM LatA for 30 minutes. **c,** Correlation heatmaps for each granule membrane marker analyzed, displaying Pearson’s correlation coefficients between the effects of knockdowns on exocytosis when treated with different stimuli. The data set contained 10 CRISPRi knockdown cell lines and control cell lines with (+) or without (-) a safe targeting sgRNA. **d**-**e**, Scatter plots comparing the effects of gene knockdowns on degranulation in responses to (**d**) 100 nM fMLF with LatA versus 100 nM LTB4 with LatA and (**e**) 100 nM fMLF with LatA versus opsonized Zymosan A, corresponding to data shown in **c**. Data are normalized to the mean of control samples. Data points and error bars represent the mean and ±s.e.m. for n=3 independent biological replicates.

To determine whether the same components link each stimulus to degranulation responses, we measured exocytosis of CD35, CD11b, CD63, and CD107a in response to each stimulus for a focused panel of CRISPRi knockdown lines (Fig. 7c-e, Extended Data Fig. 6). We found that the effects of the knockdowns on degranulation in response to fMLF and LTB4 were tightly correlated, with knockdown of *GNA15* strongly impairing the response in both cases (Fig. 7c,d). For CD35 and CD11b, we noticed that the apparent surface levels typically declined slightly in response to zymosan stimulation, likely because these two receptors bind to the opsonized target. The change in surface levels of these receptors may represent competing effects of secretion and binding to the target. Even so, the effects of gene perturbations on CD35 surface level changes were correlated for all three stimuli, and the effects on CD11b surface level changes were moderately correlated. In contrast, the effects of knockdowns on CD63 exocytosis and CD107a exocytosis were largely uncorrelated between the GPCR inputs and the opsonized zymosan input (Fig. 7c,e). Furthermore, knockdown of *GNA15* had no significant effect on zymosan-induced degranulation (Fig. 7e). Our results indicate that Gα15 promotes degranulation downstream of at least two chemoattractant GPCRs, but its role is likely specific to GPCR pathways.

### G*α*15 is necessary for sustaining elevated cytosolic Ca2+ concentrations

Gα15 is in the Gq family of G-proteins, and it signals through PLCβ to promote calcium signaling. We reasoned that Gα15 may contribute to degranulation by controlling calcium responses downstream of FPR1, even though FPR1 primarily signals through Gαi. First, we confirmed the degranulation phenotype for *GNA15* by comparing it to phenotypes for *GNAI2* and *RAB27A* using CRISPRi knockdown cell lines in small-scale experiments. We confirmed gene knockdown for each of these lines using qPCR (Extended Data Fig. 7a-c). Consistent with the results of our screens, knockdown of each gene caused defects in degranulation as measured by both CD107a and CD63 surface markers (Fig. 8a,b, Extended Data Fig. 7d,e). Also consistent with our screens, knockdown of *GNA15* caused a stronger defect in degranulation than knockdown of *GNAI2*.

**Fig. 8:**
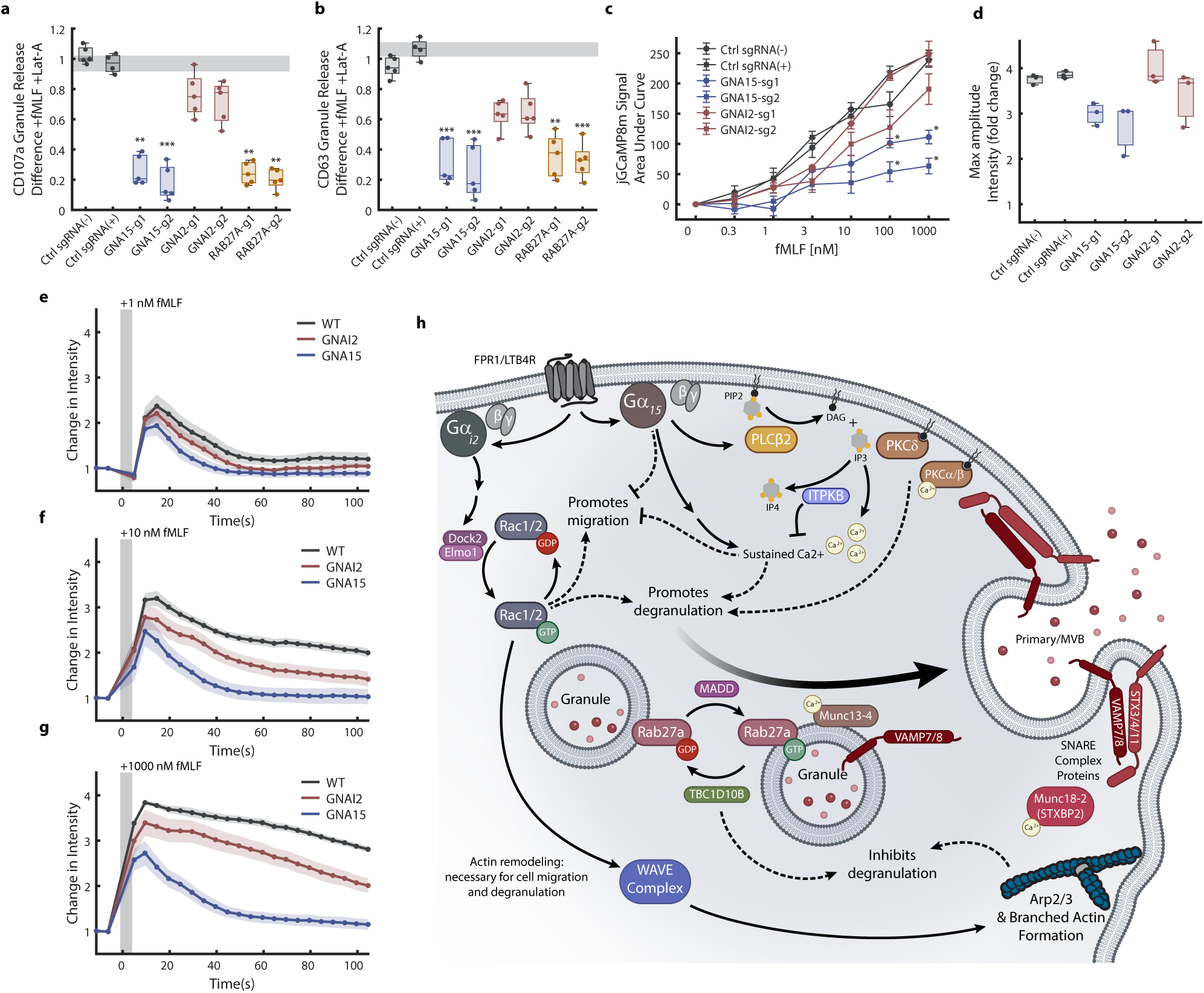
GNA15 is necessary for sustained intracellular Ca2+ signaling. **a**-**b**, Boxplots quantifying the effect of CRISPRi knockdown of GNA15, GNAI2, or RAB27A on degranulation in response to 100 nM fMLF after pretreatment with LatA, measured by the difference in cell surface expression of CD107a (**a**) or CD63 (**b**). Data normalized to the mean of control samples. Dots indicate data from n=5 independent biological replicates. P-values were calculated for each condition against the control sgRNA(+) condition using a nonparametric permutation approach with a Bonferonni correction for multiple comparisons. **c-g**, Calcium signaling responses of control dHL-60 cells or cells with CRISPRi knockdown of GNAI2 or GNA15 were measured in response to different doses of fMLF using a genetically integrated jGCaMP8m calcium sensor. (**c**) Dose-response measuring the difference in the integrated calcium response, calculated as the area under the curve for the response timecourse, comparing control and knockdown cell lines. Error bars represent the s.e.m. for n=3 independent biological replicates. P-values were computed using Welch’s ANOVA with a Games-Howell multiple comparisons test for comparisons between knockdown lines and the sgRNA(+) control cell line for each fMLF dose. (**d**) Comparison of the maximum amplitude of the calcium response for the experiments with a 1000 nM fMLF stimulus dose. Dots represent independent biological replicates. Timecourses of responses to (**e**) 1 nM fMLF, (**f**) 10 nM fMLF, and (**g**) 1000 nM fMLF are shown. Data points and curves represent the mean response, and the shaded error regions represent the s.e.m. values. The gray shaded region indicates the time of stimulus addition. **h**, Schematic diagram of the pathways downstream of chemoattractant GPCRs controlling the neutrophil degranulation response. The schematic includes roles of components identified and characterized in this study, including Gα15, the WAVE complex, MADD, TBC1D10B, ITPKB, and specific isoforms of protein kinase C.

To test the calcium signaling hypothesis, we then made CRISPRi knockdown cell lines for *GNA15* and *GNAI2* in cells that also express the jGCaMP8m genetically encoded fluorescent sensor for cytosolic calcium^55^. For each gene, we made two independent cell lines using different sgRNAs. We wanted to compare calcium signaling phenotypes for *GNA15* to the phenotype of a *GNAI2* knockdown, since Gαi represents an alternative pathway to generate calcium signals downstream of FPR1, through Gβγ-mediated activation of PLCβ^56^.

We then measured the calcium responses in control cells and cells with *GNAI2* or *GNA15* knocked down. We found that knockdown of GNAI2 caused only a modest reduction in the calcium response (Fig. 8c-g), with only the second sgRNA resulting in apparent defects in the maximum response amplitude (Fig. 8c) and the response at 60 s after stimulation (Fig. 8d). In contrast, knockdown of *GNA15* caused a partial reduction in the maximum response amplitude, but an almost complete loss of the response at 60 s and longer after stimulation (Fig. 8c-g). While a modestly reduced calcium response was apparent with a low 1 nM fMLF stimulus, the effect of the *GNA15* knockdown was stronger and more obvious at higher stimulus concentrations (Fig. 8d-g). These results indicate that Gα15 promotes stronger and sustained calcium signaling responses downstream of FPR1, particularly for the higher input doses necessary to trigger efficient degranulation.

## Discussion

In this work, we completed genome-wide screens to identify genes promoting and limiting degranulation in a neutrophil model. We did this by taking advantage of the large fold-change in the surface exposure of a granule membrane protein to apply highly efficient magnetic cell separation. We were able to separate approximately 400 million cells into populations heavily enriched for degranulating and non-degranulating subsets in a matter of minutes, providing strong statistics for downstream analysis. Our screens efficiently identified known components of the neutrophil degranulation pathway, as well as many new candidates, updating our understanding of this pathway and its relationship to the chemotaxis pathway (Fig. 8e). Our strategy is also highly adaptable, and could be applied to a range of regulated secretion processes.

One goal of our study was to get insights into the factors limiting degranulation in the absence of LatA treatment. Prior work implicated the mesh of membrane-associated cortical actin as a barrier that limits granule secretion in multiple cell types^25,27,44,57,58^. However, the cortical actin network is complex, and our results provide insight into the components and processes that play the largest roles. Our identification of branched actin assembly as a dominant process limiting secretion was consistent with the known role of Arp2/3 in assembling cortical actin. However, it was somewhat surprising that the WAVE complex components were equally strong hits, as this complex is largely associated with dynamic actin assembly in protrusive regions, as opposed to the bulk of the cortex. We did not identify other Arp2/3 activators as hits, suggesting that branched actin generated by the WAVE complex plays a particularly important role in limiting degranulation, although it remains unclear if this role is through protrusive activity or through generating a bulk of cortical actin.

A second major goal of our study was to identify signaling components that distinguish the control of degranulation from chemotaxis downstream of attractant GPCRs. In this regard, we identified Gα15 as a potential pathway branch point. Loss of Gα15 severely limits degranulation, but does not impair chemotaxis. Molecularly, we found that Gα15 promotes stronger and more sustained calcium signaling downstream of receptor activation, which may be important for driving calcium-dependent steps, such granule fusion with the plasma membrane. The role of Gα15 appears to be specific to GPCR inputs, but our results with LTB4 and fMLF indicate that it may play a shared degranulation-promoting role downstream of multiple chemoattractant GPCRs. As attractant GPCRs differ in their potency for triggering degranulation, it is possible that variability in the strength of their coupling to Gα15 underlies differences in potency.

Our results indicate that both Gαi (*GNAI2*) and Gα15 are needed for efficient degranulation. The role of Gαi is consistent with prior studies demonstrating that Gαi is needed for almost all responses to fMLF inputs, including degranulation^16,59^. In particular, treatment of neutrophils with pertussis toxin, which inactivates Gαi G-proteins, blocks chemotaxis, signaling responses including calcium influxes, and degranulation. These results raise a question of how the two G alpha subunits work together to coordinate the degranulation response. A simple possible explanation is that Gαi activates downstream effectors such as DOCK2 and Rac2 that are shared between the chemotaxis and degranulation pathways, and that Gα15 provides the complementary strong calcium signals needed to trigger degranulation. However, it is also possible that Gαi plays a role in promoting receptor coupling to Gα15, and that each G-protein may have additional effectors that contribute.

While our study was focused on neutrophil degranulation, this process shares close mechanistic connections with other regulated secretion processes, including degranulation by cytotoxic lymphocytes, insulin secretion by pancreatic beta cells, and the release of exosomes. Indeed, there is considerable overlap between the strongest hits in our screens and the components known to be required for the secretion of lytic granules by T and NK cells^19,28^. Furthermore, the release of exosomes by neutrophils has been implicated in modulating inflammatory responses in multiple contexts^40,60,61^. Exosomes are released when multivesicular bodies fuse with the plasma membrane. Rab27 proteins regulate this fusion event and CD107a is found on multivesicular bodies^29,40^. Thus, secretion of exosomes by neutrophils could be closely mechanistically linked to degranulation. More broadly, many of the components that we identified as regulators of degranulation in dHL-60 cells are strong candidates to control neutrophil exosome release and Rab27-regulated secretion events in other cell types.

## Methods

### Degranulation and Differentiation Genome-Wide CRISPR-Cas9 Screens

To screen for regulators of degranulation, magnetic particles were used to physically separate degranulating cells from non-degranulating cells. Each population was then sequenced to quantify guide enrichment and call hits. The cell line used, HL-60, was differentiated into a neutrophil-like state prior to performing degranulation screens. To identify regulators of this differentiation process, cell populations were sequenced before and after differentiation.

#### Cell preparation

HL-60 cells expressing an exogenous copy of FPR1 and Cas9 were prepared with a pooled library of sgRNA plasmids. A pre-differentiation sample of 5.0 x 10^7^ cells was frozen. To prepare cells for degranulation screens and obtain a sample of differentiated cells, 2.5 x 10^8^ HL-60 cells were cultured at an initial density of 2.0 x 10^5^ cells/ml in 1,250 ml of RPMI +2% nutridoma and 1.3% DMSO. The cells were incubated for 6 days to reach a differentiated state (dHL-60). At day 6, the cells had proliferated to a total of 1.2 x 10^9^ cells, which were divided into three separate pools of 4.0 x 10^8^ cells in 10 ml of RPMI +10% FBS each for the screens. One pool of cells was frozen for use in the differentiation screen. The remaining cells were used for an fMLF stimulus screen and an fMLF and Latrunculin-A stimulus screen.

#### Magnetic particle preparation

Smooth surface, 3.5 um Streptavidin Coated Magnetic Particles (Spherotech catalog # SVMS-30-10) were washed with 1X PBS and resuspended at 10 mg/ml in 1X PBS. Biotin anti-human CD107a (LAMP-1) antibody (Biolegend catalog # 328604) was added at a ratio of 5 ug antibody per 1 mg magnetic particle. The mixture was incubated with rotation for 30 minutes at room temperature then washed twice with 1X PBS to remove excess antibody.

#### Stimulation and separation

Cells were incubated on a 37C heat block before and during stimulation. For the fMLF and Latrunculin-A treated sample, 1 uM Latrunculin was added and the cells were mixed by gentle inversion. Next, 1 uM fMLF (Sigma-Aldrich catalog # 47729) was added to both samples, mixed, and allowed to incubate for 3 minutes. The cells were immediately placed on ice following stimulus and incubation, then centrifuged for 5 min at 200 rcf and 4C. The cells were resuspended in 10 ml FACS buffer containing magnetic particles and rotated at 4C for 25 minutes. To separate the cell populations, the tubes were placed on magnets to apply a uniform magnetic field that pulled cells with magnetic particles bound to them to the side of the tube while unbound (non-degranulating cells) remained in solution. After 5 minutes of incubation, the tube was inverted, pouring out the unbound cells to a separate tube. This process was repeated twice to ensure the cell populations were thoroughly separated. Next, the collected cells were suspended in RPMI containing 20% FBS and 10% DMSO and stored at −80C until DNA isolation for sequencing.

### Analysis of genome-wide screens

To identify guide RNAs and genes significantly enriched or disenriched in cell populations of our screens, we used the log-likelihood scoring system, effect size estimation, and empirical p-value computation strategy that we previously described^33^. The key elements of the approach are summarized here for clarity.

#### Computation of CLO scores at the guide and gene levels

The CRISPR Log Odds (CLO) scores provide a measure of the enrichment of an sgRNA or a gene, accounting for empirical measurement noise through a negative binomial model in which measurement variance is estimated as a function of mean read number. Scores for individual sgRNAs are computed first, and then guide-level scores are summed to obtain gene-level scores. The formulas for the guide and gene-level CLO scores are given by:

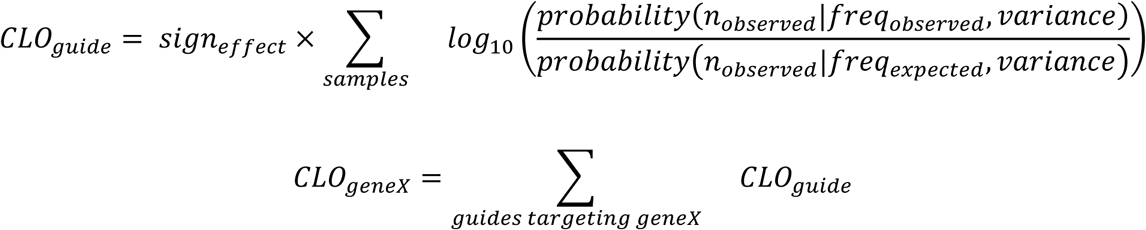

The sign (positive or negative) indicates the direction of the effect (enriched vs depleted). The probabilities of obtaining the observed number of reads for a sample are computed using negative binomial models parametrized by their mean and variance, where the mean is equal to the observed number of reads (observed) or the observed guide frequency across all samples multiplied by the effective total number of reads for that sample (expected). The effective total number is computed using only the abundance of control guides, to account for potential distortions due to large-scale enrichment or depletion of a subset of gene-targeting guides. The variance is estimated by fitting the variance for control guides using a linear fit of the log variance as a function of the log mean number of reads.

#### Computing p-values

Guide-level empirical p-values were computed for each guide using its rank order relative to the scores of all control guides. P-values assessing guides as potential hits in the negative direction were computed as the number of control guides with a lower CLO score plus 0.5, divided by the total number of control guides. P-values assessing guides as potential hits in the positive direction were computed as one minus the p-value for the negative direction. Gene-level p-values (for each direction) were computed using the algorithm for combining p-values from Bailey and Gribskov^62^ based on the null model that the p-values come from a uniform distribution.

#### Computing effect size estimates for individual guides

Raw effect sizes for each guide were computed as the log_2_ fold-change in read frequency between conditions. To account for measurement noise which can result in large apparent effect sizes for guides with small read numbers, a Bayesian maximum likelihood approach was used. A smoothed distribution of the raw effect sizes (restricted to guides with an average of at least 100 reads per sample) was used for the prior distribution. Posterior likelihoods were computed over a grid of potential “true” guide frequencies for each sample. The final estimated effect size was taken to be the log_2_ fold-change between the frequencies with the highest posterior likelihoods.

### Cell culture

All cell lines were maintained in New Brunswick Galaxy S Series 170 S CO_2_ incubators (New Brunswick Scientific, Edison, NJ) at 37°C with 5% supplemental CO_2_. For imaging experiments, a “modified L-15” imaging media (Leibovitz’s L-15 media lacking dye, riboflavin, and folic acid) (UC Davis Biological Media Services) was used to minimize media autofluorescence and to maintain media pH without supplemental CO_2_. Cell lines were regularly tested for mycoplasma. No mycoplasma contamination was detected for any cell line used in this work.

#### HL-60 cell culture

HL-60 cells were cultured in RPMI-1640 (Gibco, catalog #72-400-120) media with 9% heat-inactivated fetal bovine serum (FBS) (Sigma-Aldrich, catalog #F4135). The cell line used for all experiments was a subline of HL-60 cells known as PLB-985. We previously validated the genetic identity of these cells by SNP analysis^24^. The cell line used was originally obtained as a gift from Dr. Orion Weiner (University of California, San Francisco). Cells were passaged every two to three days and maintained at a culture density between 1.0 × 10^5^ and 2.0 × 10^6^ cells/ml. Cells were differentiated into a neutrophil-like state by culturing at an initial density of 2 × 10^5^ cells/ml in RPMI-1640 with 5% heat-inactivated FBS, 100 mg/ml, 1.3% DMSO, and 2% Nutridoma-CS (Roche, catalog #11363743001). Cells were incubated for 6 days before use in experiments, at which point they were referred to as dHL-60 cells.

#### HEK-293T cell culture

HEK-293T cells (ATCC CRL-11268) were used for lentiviral production. Cells were cultured in high glucose Dulbecco’s modified Eagle’s medium (Sigma-Aldrich, D5671) that was supplemented with 9% FBS and 1% Glutamax (Gibco, catalog # 35050061). Cells were passaged at confluency and detached from culture dishes using TrypLE Express (Gibco, catalog # 12605036).

### Cell-line generation

HL-60 cell lines containing exogenously expressed genes were generated using 2nd generation lentiviral-mediated gene transfer. The packaging vector used was a gift from Dr. Lifeng Xu (pMD.G, University of California, Davis), while the envelope vector was a gift from Dr. Peter Lewis (pCMV-dR8.2, University of Wisconsin, Madison). To create the lentivirus, HEK-293T cells were transfected in Opti-MEM (Gibco, catalog # 51985-034) using TransIT-2020 transfection reagent (VWR, catalog # 10767-014). Media was changed to DMEM containing 10% FBS after 12 hours. Supernatant was collected at 24 and 48 hours post media change and filtered (PES 0.45µm; catalog # 25-246). Lentivirus containing supernatant was concentrated 50-fold as described by the manufacturer (Origene, catalog # TR30026), flash frozen in liquid nitrogen and stored at −80°C. HL-60 cells were treated with 50 μl of the concentrated lentivirus and 2 mg/ml Polybrene (Sigma; catalog # H9268-5G). For genetic constructs with antibiotic resistance, infected cells were centrifuged (200g, 3 minutes) and resuspended in selection media (supplemented RPMI-1640 with antibiotics; 500 µg/ml Hygromycin, 5 µg/ml Puromycin, or 50 mg/ml Neomycin) approximately 24 hours post lentiviral infection.

#### Individual sgRNA knockdown cell lines

To efficiently create all knockdown cell lines, the lenti virus production and subsequent transduction was adjusted slightly. HEK-293T cells were transfected with polyethylenimine, (Polysciences, Catalog # 223966) in high glucose Dulbecco’s modified Eagle’s medium (Sigma-Aldrich, D5671) supplemented with 9% FBS and 1% Glutamax. Supernatant was collected 48 hours after transfection and filtered. Lentivirus containing supernatant was concentrated 50-fold as described by the manufacturer (Origene, catalog # TR30026), flash frozen in liquid nitrogen and stored at −80°C. HL-60 cells were treated with 10 μl of the concentrated lentivirus before selection for genetic integration with 5 µg/ml Puromycin.

### Human primary neutrophil isolation

Ethical approval for the study of neutrophils from adult healthy controls was granted by the Institutional Review Board from the University of California, Davis (IORG0000251). Participants gave written, informed consent. Blood was collected by finger prick using a microlet lancing device. Neutrophils were isolated using negative selection with the EasySep Direct Human Neutrophil isolation kit (STEMCELL Technologies, catalog no. 19666) following the manufacturer’s instructions.

### Flow-cytometry

Flow cytometry was used with fluorophore conjugated antibodies to quantify cell surface expression of receptors and granule membrane proteins. Live cells were identified using gates based on FSC and SSC parameters. Cells were collected and, to remain consistent across experiments, were transferred to modified L-15 containing 10% FBS. Cells were incubated on 37C heat blocks during stimulation. Following stimulation, cells were transferred to antibody staining buffer (0.5% bovine serum albumin and 0.05% sodium azide in PBS) and incubated on ice for 30 minutes. After fluorescent labeling, the samples were washed with ice-cold FACS buffer and kept on ice until fluorescence was quantified via flow cytometry. Channel cross-talk was determined by acquiring samples with individual antibody stains. Flow cytometry data was analyzed using custom MATLAB scripts.

#### Degranulation assay

2 x 10^5^ cells in 500 ul L-15 +10% FBS for each condition were transferred to 1.5 ml tubes and on a 37C heatblock. Conditions with chemical inhibitors were pretreated for 30 minutes at 37C. Stimulation was applied and tubes were inverted to mix rapidly. Samples were incubated for 3 minutes and then transferred to ice to slow cellular responses. Cells were then transferred to antibody staining buffer (0.5% bovine serum albumin and 0.05% sodium azide in PBS) and incubated on ice for 30 minutes. Staining conditions were a combination of the following antibodies: 1:100 FITC anti-human CD35 (Biolegend; catalog # 333404), 1:100 AlexaFluor-647 anti-human CD107a (Biolegend; catalog # 328612), and 1:100 Pe-Cy7 anti-human CD63 (Biolegend; catalog # 353010) Channel crosstalk compensation was not needed. Data was acquired on a BD FACS Canto II flow cytometer: 2 laser (Blue, Red), 6-color (BD Biosciences, Franklin Lakes, NJ).

#### High throughput degranulation assay

2 x 10^5^ cells in 100 ul L-15 +10% FBS for each condition were transferred to 96-well V-bottom plates. Cells were incubated at 37C in a benchtop incubator (Eppendorf; ThermoMixer C). Conditions with chemical inhibitors were pretreated for 30 minutes at 37C. Stimulation was applied in 2X concentrations for rapid mixing using multichannel pipettes, incubated for 3 minutes, and then transferred to ice to slow cellular responses. Cells were then transferred to antibody staining buffer (0.5% bovine serum albumin and 0.05% sodium azide in PBS) and incubated on ice for 30 minutes. Staining conditions were a combination of the following antibodies: 1:100 Brilliant Violet 421 anti-human CD35 Antibody (Biolegend; catalog #333416), 1:100 Alexa Fluor 488 anti-human CD63 Antibody (Biolegend; catalog #353038), 1:25 APC anti-human FPR1 Antibody (Biolegend; catalog #391610), 1:100 Alexa Fluor 700 anti-human CD11b Antibody (Biolegend; catalog #301356) and 1:100 PE/Cyanine7 anti-human CD107a (Biolegend; catalog #328618). Channel crosstalk was corrected with a compensation matrix (Extended Data Fig. 5b). Data was acquired on a BC Cytoflex flow cytometer: 4 laser (Violet, Blue,Yellow, Red), 13-color, 96-well plate acquisition (Beckman Coulter, Indianapolis, IN).

#### Human primary neutrophil degranulation assay

2 x 10^5^ cells primary neutrophils were resuspended in 1.5ml tubes and incubated with corresponding chemical inhibitors in L-15 +1% BSA for 30 minutes at 37C. Stimulation was applied in 2X concentrations for rapid mixing, incubated for 3 minutes, and then transferred to ice to slow cellular responses. Cells were then transferred to antibody staining buffer (1% BSA and 0.05% sodium azide in PBS) and incubated on ice for 30 minutes. Cells were stained with 1:100 Alexa Fluor 488 anti-human CD63 Antibody (Biolegend; catalog #353038), 1:25 APC anti-human FPR1 Antibody (Biolegend; catalog #391610) and 1:100 PE/Cyanine7 anti-human CD107a (Biolegend; catalog #328618).

### Chemical inhibitors

Chemical inhibitors used included the actin polymerization inhibitor Latrunculin A (Millipore Sigma; catalog #428026); Arp2/3 Complex inhibitors CK-666 (MedChemExpress; catalog #HY-16926) and CK-869 (MedChemExpress; catalog #HY-16927); SLK/LOK inhibitor SLK/STK10-in-1 (MedChemExpress; catalog #HY-132868); and formin inhibitor SMIFH2 (MedChemExpress; catalog #HY-16931). Concentrations are specified where used.

### Statistical analysis

For comparisons involving multiple experimental groups for which the data was consistent with the normality assumption, a one-way ANOVA and a Tukey’s multiple-comparison test were used to assess significance and compute pairwise p-values. A Welch’s ANOVA and the Games-Howell post hoc test were used in cases where the data was not consistent with the homogeneity of variance assumption for some samples.

For comparisons involving multiple experimental groups for which the data was not consistent with the normality assumption, a nonparametric permutation analysis was used to compute pairwise p-values, and the p-values were corrected using the Bonferroni procedure to control the familywise type I error rate. A null distribution containing 10^6^ permutations was generated using pooled data from all experimental groups in the corresponding figure panel. P-values were calculated using the Monte Carlo approach, where the absolute value of the difference between sample means was compared to the frequency of an equal or greater absolute difference within the null distribution.

The Shapiro-Wilk’s test and Levene’s test were performed on residuals to test normality and homogeneity of variance assumptions.

### Under agarose Chemotaxis

Cell migration and chemotaxis were measured using an under agarose assay based on photo-uncaging of the chemoattractant fMLF with small modifications from our previously described approach^49^. Key details are described here for clarity. Glass bottom 96-well plates (Cellvis, #P96-1.5H-N) were coated with 1% bovine serum albumin and dried at 65°C to prevent cell over-adhesion. Cells were labeled with SybrSafe (ThermoFisher, #S33102) at a 1:30,000 dilution to allow fluorescent imaging of nuclei for cell tracking. Approximately 1000 dHL-60 cells with the indicated CRISPRi constructs were plated in the center of each well in a 5 μL volume. Cells were allowed to adhere for 5 minutes, and then 195 μL of a liquid media-agarose mixture (1.5% low melting temperature agarose (Bio Basic Inc, #AB0015) in modified L15 media with 5% FBS and 100 nM N-nitroveratryl fMLF (Nv-fMLF) at 37C) was added on top. The agarose was allowed to solidify for 40 minutes at room temperature, and then moved to the heated microscope chamber at 37 C. The plate was allowed to equilibrate to 37C for 40 minutes before imaging. Cells were imaged every 30s at 4x magnification in the YFP channel (combined emission bandpass of ∼525-550 nm) with a Nikon Ti2 inverted microscope to allow cell tracking. After 7.5 minutes, a chemoattractant gradient was generated using UV photo-uncaging of Nv-fMLF. Uncaging was performed with 0.5 s of patterned UV illumination using a Lumencor Aura III with a 360/28 nm bandpass filter and Polygon1000-G digital micromirror device (Mightex). The uncaging region was a disc of approximately 1 mm radius in the center of the field of view. Imaging was continued for 17.5 minutes with continued uncaging pulses of 0.5 s every 30 s to maintain the attractant gradient.

### Cell tracking and statistics

For the chemotaxis experiments, cell nuclei were segmented and tracked using previously developed custom MATLAB code^49^. Briefly, cell nuclei were segmented in each frame and tracked between frames to create trajectories. Movement vectors between successive frames were created from trajectories. Cell speed before and after gradient generation were computed over the corresponding time periods as the mean length of movement vectors divided by the time interval of 0.5 minutes. Directed cell speed was computed as the mean length of the projection of the movement vector onto the gradient direction divided by the time interval of 0.5 minutes. Movement angles were computed as the angle between the movement vector and the direction towards the center of the gradient. Mean angular bias was computed, considering only movement vectors of length 2.5 pixels or greater, as the mean of 90 minus the movement angle. Since a mean movement angle of 90 degrees is expected for random movement, this measures mean bias in degrees towards the gradient center. Phenotype scores were computed by taking raw measurements, subtracting the mean value of the three control cell lines, and dividing by this same mean. Thus, a phenotype score of zero represents no difference from the control, and nonzero scores indicate a fractional difference relative to the control. For the scatter plots in Figure 6b and 6c, shaded error regions were computed by empirically fitting the measurement variability across replicates as a sum of two gaussian distributions^49^. The shaded region represents the predicted range encompassing 99% of averaged scores for a gene with zero phenotype (no effect).

### Plasmid creation

Plasmids were amplified by transforming One Shot Stbl3 chemically competent E. Coli (Invitrogen, catalog no. C737303). E. Coli were grown at 37°C in Luria Broth containing antibiotics. All isolated plasmid sequences were verified by Sanger Sequencing.

#### sgRNA knockdown plasmids

Lentiviral expression vector was linearized using restriction enzymes *BlpI* (Thermo Fisher Scientific, Catalog # FD0094) and *BstXI* (Thermo Fisher Scientific, Catalog # FD1024). Forward and reverse DNA oligo sequences containing the sgRNA guide sequence were annealed and phosphorylated using T4 PNK (New England Biolabs, Catalog # M0201S). Annealed oligo sequences were ligated into the linearized vector plasmid using Quick Ligase (New England Biolabs, Catalog # M2200S). The oligo sequences used to make these sgRNA plasmids are included in Supplementary Table 1.

#### jGCaMP8m cloning

jGCaMP8m DNA was acquired from Addgene (Plasmid #162372). The jGCaMP8m sequence and a neomycin resistance gene sequence were amplified by PCR and incorporated into a linearized lentiviral expression vector by Gibson Assembly.

### Detection of gene expression using two-step RT-qPCR

Total RNA was extracted from dHL-60 cells on 4 different days. Total RNA extraction was done using Qiagen RNeasy RNA extraction kit according to the manufacturer’s protocol. 1000 ng of purified messenger RNA (mRNA) was used to reverse transcribe to complementary DNA (cDNA) using the 5× All-In-One RT Master Mix kit (abm). Around 20 ng cDNA template was used in the RT-PCR reaction (KiCqStart® SYBR® Green qPCR ReadyMix™) with three technical replicates for each condition. 4 rounds of qPCR were performed using cDNA obtained from cells differentiated on different days. Conditions used in the LightCycler® 480 Instrument were: 95 for 5 s, 45 cycles of 95 °C for 10 s, 55 °C for 15 s, 72 °C for 20 s, Melting curve of 95 °C for 5 s, 65 °C for 1 m, Cooling. Primers were designed to span exon-exon junctions for target genes. Primer sequences are listed in Supplementary Table 1. To quantify enrichment in gene expression in targeting guide-expressing cells relative to the safe- or non-targeting guide-expressing cells, we used the delta-delta Ct (ΔΔCt) method.

### Intracellular Ca2+ kinetics

Intracellular Ca^2+^ kinetics were measured by imaging live cells that expressed the genetically encoded calcium sensor, jGCaMP8m. dHL-60 cells were centrifuged at 200 g for 3 min at room temperature and resuspended in warmed modified L-15 media containing 2% FBS at a density of 5.0×10^5^ cells/ml. Cells (5.0×10^4^ cells/per well) were plated in 96-well optical-glass bottomed imaging plates (Cellvis, catalog no. P96-1.5H-N) coated with polyD-lysine (“PDL,” Sigma-Aldrich, catalog no. P6407) to minimize cell washout. To apply the coating, PDL (200 mg/ml) was added to each well and incubated at room temperature for 30 min. After incubation, wells were washed twice with an excess of deionized water and dried thoroughly at 65°C for at least 30 min. Image frames were acquired approximately every 2 seconds. During image acquisition cells were stimulated by pipetting 2X chemoattractant solutions into the imaging wells. Quantification of fluorescent signal was performed using custom MATLAB scripts. Briefly, our image processing workflow included removal of samples with pipetting artifacts, a flat field correction, background subtraction and object detection, followed by calculation of the mean fluorescence of the cell occupied area in an image frame. To account for day-to-day variability, data were normalized to the mean of the two image frames prior to stimulation. Images were acquired on a Nikon Ti2 inverted microscope.

## Acknowledgments

This work was supported by the NIH under the grants DP2HD094656 and R01GM148769 to SRC, the Floyd and Mary Schwall Dissertation Year Fellowship, Lurie Graduate Student Award, and National Institutes of Health training grant T32GM007377 to SML, by the UC Davis Dissertation Year Fellowship to EA, and the University of California, Davis. We thank Bridget McLaughlin and Jonathan Van Dyke for their technical assistance at the University of California Davis Flow Cytometry Shared Resource Laboratory with funding from the NCI P30 CA093373 (Comprehensive Cancer Center) and S10 OD018223. We thank current Collins lab members and alumni, as well as Prof. Nicholas Marsh-Armstrong, Prof. Karen Zito, and Prof. Frances Mercer for valuable discussions and their input on this project. We thank Jonathan Potter and Kristi Bispo for technical assistance and discussions. We thank the lab of Jonathan Weissman (Broad Institute) for sharing reagents and protocols.

The authors declare no competing financial interests.

## Extended Figure Legends

**Extended Data Fig. 1: Validation of Cas9 function and relationship between screen hits and corresponding gene expression levels**. **a**, GFP expression levels were measured by flow cytometry for dHL-60 cells with stable integration of a Cas9 expression construct and/or a construct expressing GFP and an sgRNA that targets GFP. The reduction of GFP positive cells when both constructs are integrated serves as a measure of Cas9 activity. **b**, Histogram of gene expression levels in log_10_ transcripts per million (TPM) in dHL-60 cells (data from Rincon et al^24^). **c**, Frequency of screen hits (defined as passing the 5% FDR threshold) as a function of gene expression level for each of the three genome-wide screens. Gene expression values were binned and the fraction of genes passing the 5% threshold was calculated for each bin. **d-f**, Scatter plots of gene expression level versus the genome-wide screen logP score for the differentiation (**d**), fMLF only degranulation (**e**), and fMLF plus LatA degranulation (**f**) screens. The gray shaded region indicates the logP score range falling below the 5% FDR threshold for each screen. Genes with no detected transcripts are shown at −4 on the gene expression axis.

**Extended Data Fig. 2: Continuation of overview of hits from genome-wide degranulation screens.** The schematic is a continuation of the data shown in Fig. 3 and shows logP score heatmaps for genes, organized by functional categories. All genes shown passed a 5% FDR threshold for at least one degranulation screen. Hits for the differentiation screen were excluded. For each gene, the score for the fMLF only screen is shown on the left, and the score for the screen with fMLF and LatA is shown on the right. Functional categories were assigned manually based on prior literature.

**Extended Data Fig. 3: Granule membrane protein distribution across granule subtypes and measured changes in surface expression in response to fMLF. a-d.** Relative abundance of granule marker proteins across granule subtypes. Raw data was obtained from Rørvig et at., 2013^26^ based on mass spectrometry measurement of protein abundances in density gradient-separated neutrophil granule fractions. **e**-**g**. Representative histograms showing cell surface levels of the granule membrane proteins CD63, CD35, and CD11b in dHL-60 cells following stimulation with the indicated concentrations fMLF. **h**-**j**. Representative histograms showing cell surface levels of the granule membrane proteins CD63, CD35, or CD11b in dHL-60 cells following treatment with the indicated concentrations of fMLF after pretreatment with LatA for 30 minutes.

**Extended Data Fig. 4: Analysis of degranulation hits across granule subtypes. a, b**. Heatmaps comparing the effects of CRISPRi knockdowns of the selected genes on exocytosis of different granule subtypes in dHL-60s. The effects were measured as the change in cell surface expression of the indicated granule membrane proteins (CD35, CD11b, CD63, and CD107a) after stimulation, then normalized to the mean of the control samples. Cells were stimulated with 100 nM fMLF (**a**) or 10 nM fMLF and 1 uM LatA (**b**). **c-e**, Scatter plots from data in (**a**), comparing the effects of gene knockdowns between granule sub-type exocytosis. Data are normalized to the mean of three control samples. Data points and error bars represent the mean and ±s.e.m. for n=3 independent biological replicates.

**Extended Data Fig. 5: Measurement of cell surface FPR1 protein levels for CRISPRi knockdown cell lines**. **a**, Box plots show cell surface levels of FPR1 measured by flow cytometry (in parallel with measurements of CD35, CD63, CD107a, and CD11b) for the indicated CRISPRi gene knockdown cell lines. For each gene (except FPR1), measurements for two knockdown cell lines are shown, with sg1 on the left and sg2 on the right. These cell lines are the same cell lines used for the experiments in Fig. 4 and Extended Data Fig. 4. **b**, The compensation matrix used to correct cytometry measurements for the experiments in Fig. 4, Extended Data Fig. 4, and panel (**a**).

**Extended Data Fig. 6: Comparison of the effects of degranulation hit gene knockdowns for different stimuli.** Scatter plots comparing the effects of gene knockdowns on exocytosis when treated with different stimuli. Each scatter plot shows the data for 10 CRISPRi knockdown cell lines and control cell lines with (+) or without (-) a safe targeting sgRNA. **a-c**, Scatter plots comparing the effects of gene knockdowns on degranulation in response to 100 nM fMLF versus 100 nM LTB4. Degranulation was measured by the change in surface expression of CD35 (**a**), CD11b (**b**), or CD63 (**c**). **d-f,** Scatter plots comparing the effects of gene knockdowns on degranulation in response to 100 nM fMLF versus opsonized Zymosan A (at a ratio of 2 particles per cell) and measured by the change in surface expression of CD35 (**d**), CD11b (**e**), or CD63 (**f**). Data are normalized to the mean of control samples. Data points and error bars represent the mean and ±s.e.m. for n=3 independent biological replicates.

**Extended Data Fig. 7: Assessment of gene knockdown by qPCR and effects of gene knockdowns on degranulation in response to fMLF stimulation (without LatA)**. **a-c**, Quantitative PCR (qPCR) measurement of transcript levels for GNA15 (**a**), GNAI2 (**b**), and RAB27A (**c**) in control or CRISPRi knockdown dHL-60 cell lines. Data were normalized to the mean of the two control cell lines. Dots indicate independent biological replicates. **d,e**, Boxplots quantifying the effect of CRISPRi knockdown of GNA15, GNAI2, or RAB27A on degranulation in response to 100 nM fMLF (without LatA), measured by the difference in cell surface expression of CD107a (**d**) or CD63 (**e**). Data was normalized to the mean of control samples. Dots indicate data from n=5 independent biological replicates. P-values were calculated for each condition against the control sgRNA(+) condition using a nonparametric permutation approach with a Bonferonni correction for multiple comparisons.

## Supplementary Data

Supplementary Data 1: Guide level screen scores

Supplementary Data 2: Gene level screen scores

Supplementary Data 3: Degranulation screen hits and annotations

Supplementary Data 4: Chemotaxis results

Supplementary Table 1: Oligo sequences for plasmid construction and qPCR analysis

## References

1. Burgoyne, R. D. & Morgan, A. Secretory granule exocytosis. Physiol. Rev. 83, 581–632 (2003).

2. Mok, A. CH., Mody, C. H. & Li, S. S. Immune Cell Degranulation in Fungal Host Defence. J. Fungi 7, 484 (2021).

3. Lacy, P. et al. Mechanisms of Degranulation in Neutrophils. Allergy Asthma Clin. Immunol. 2, 98 (2006).

4. Kolaczkowska, E. & Kubes, P. Neutrophil recruitment and function in health and inflammation. Nat. Rev. Immunol. 13, 159–175 (2013).

5. Ley, K., et al. Neutrophils: New insights and open questions. Sci. Immunol. 3, eaat4579 (2018).

6. Lämmermann, T. & Kastenmüller, W. Concepts of GPCR-controlled navigation in the immune system. Immunol. Rev. 289, 205–231 (2019).

7. Dallegri, F. & Ottonello, L. Tissue injury in neutrophilic inflammation. Inflamm. Res. 46, 382–391 (1997).

8. Borregaard, N. & Cowland, J. B. Granules of the Human Neutrophilic Polymorphonuclear Leukocyte. Blood 89, 3503–3521 (1997).

9. Hsieh, L. L. et al. SARS-CoV-2 induces neutrophil degranulation and differentiation into myeloid-derived suppressor cells associated with severe COVID-19. Sci. Transl. Med. 17, eadn7527 (2025).

10. Matson, S. M. et al. Neutrophil degranulation in the lung microenvironment linked to idiopathic pulmonary fibrosis severity and survival. iScience 29, 114125 (2026).

11. Zhang, N., Aiyasiding, X., Li, W., Liao, H. & Tang, Q. Neutrophil degranulation and myocardial infarction. Cell Commun. Signal. 20, 50 (2022).

12. Fujishima, S. et al. Neutrophil elastase and systemic inflammatory response syndrome in the initiation and development of acute lung injury among critically ill patients. Biomed. Pharmacother. 62, 333–338 (2008).

13. Li, Y. et al. Targeting formyl peptide receptor 1 reduces brain inflammation and neurodegeneration. Science 390, eadq1177 (2025).

14. He, R., Chen, Y. & Cai, Q. The role of the LTB4-BLT1 axis in health and disease. Pharmacol. Res. 158, 104857 (2020).

15. Zhou, H. L. et al. Association between leukotriene B4-induced phospholipase D activation and degranulation of human neutrophils. Biochem. Pharmacol. 46, 139–148 (1993).

16. Becker, E. L. et al. The inhibition of neutrophil granule enzyme secretion and chemotaxis by pertussis toxin. J. Cell Biol. 100, 1641–1646 (1985).

17. Badu-Nkansah, K. A. & Collins, S. R. Leukocyte Chemotaxis and the Elusive Negative Signals that Make It Work. Cold Spring Harb. Perspect. Biol. a041792 (2026) doi:10.1101/cshperspect.a041792.

18. Yin, C. & Heit, B. Armed for destruction: formation, function and trafficking of neutrophil granules. Cell Tissue Res. 371, 455–471 (2018).

19. Kabanova, A., Zurli, V. & Baldari, C. T. Signals Controlling Lytic Granule Polarization at the Cytotoxic Immune Synapse. Front. Immunol. 9, 307 (2018).

20. Lyubchenko, T. A., Wurth, G. A. & Zweifach, A. Role of Calcium Influx in Cytotoxic T Lymphocyte Lytic Granule Exocytosis during Target Cell Killing. Immunity 15, 847–859 (2001).

21. Herrero-Turrión, M. J., Calafat, J., Janssen, H., Fukuda, M. & Mollinedo, F. Rab27a Regulates Exocytosis of Tertiary and Specific Granules in Human Neutrophils. J. Immunol. 181, 3793–3803 (2008).

22. Brzezinska, A. A. et al. The Rab27a effectors JFC1/Slp1 and munc13-4 regulate exocytosis of neutrophil granules. Traffic 9, 2151–2164 (2008).

23. Bhakta, S. B. et al. Neutrophil-like cells derived from the HL-60 cell-line as a genetically-tractable model for neutrophil degranulation. PLOS ONE 19, e0297758 (2024).

24. Rincón, E., Rocha-Gregg, B. L. & Collins, S. R. A map of gene expression in neutrophil-like cell lines. BMC Genomics 19, 1–17 (2018).

25. Jog, N. R. et al. The actin cytoskeleton regulates exocytosis of all neutrophil granule subsets. Am. J. Physiol. Cell Physiol. 292, C1690–1700 (2007).

26. Rørvig, S., Østergaard, O., Heegaard, N. H. H. & Borregaard, N. Proteome profiling of human neutrophil granule subsets, secretory vesicles, and cell membrane: correlation with transcriptome profiling of neutrophil precursors. J. Leukoc. Biol. 94, 711–721 (2013).

27. Ritter, A. T. et al. Cortical actin recovery at the immunological synapse leads to termination of lytic granule secretion in cytotoxic T lymphocytes. Proc. Natl. Acad. Sci. U. S. A. 114, E6585–E6594 (2017).

28. Medlyn, M. J. et al. MADD regulates natural killer cell degranulation through Rab27a activation. J. Cell Sci. 137, jcs261582 (2024).

29. Dahlgren, C., Carlsson, S. R., Karlsson, A., Lundqvist, H. & Sjölin, C. The lysosomal membrane glycoproteins Lamp-1 and Lamp-2 are present in mobilizable organelles, but are absent from the azurophil granules of human neutrophils. Biochem. J. 311, 667–674 (1995).

30. Mitchell, T., Lo, A., Logan, M. R., Lacy, P. & Eitzen, G. Primary granule exocytosis in human neutrophils is regulated by Rac-dependent actin remodeling. Am. J. Physiol.-Cell Physiol. 295, C1354–C1365 (2008).

31. Belliveau, N. M. et al. Whole-genome screens reveal regulators of differentiation state and context-dependent migration in human neutrophils. Nat. Commun. 14, 5770 (2023).

32. Morgens, D. W. et al. Genome-scale measurement of off-target activity using Cas9 toxicity in high-throughput screens. Nat. Commun. 8, 15178 (2017).

33. Akdoğan, E., Lundgren, S. M., Kamber, R. A., Bassik, M. C. & Collins, S. R. Parallel CRISPR screens reveal pathways controlling the cell surface levels of the attractant receptor FPR1. Commun. Biol. 9, 668 (2026).

34. Zhao, X. W. et al. Defects in neutrophil granule mobilization and bactericidal activity in familial hemophagocytic lymphohistiocytosis type 5 (FHL-5) syndrome caused by STXBP2/Munc18-2 mutations. Blood 122, 109–111 (2013).

35. Ilarraza, R. et al. Rac2 regulates immune complex–mediated granule polarization and exocytosis in neutrophils. J. Leukoc. Biol. 114, 116–125 (2023).

36. Kat, M. et al. GDP/GTP exchange factor MADD drives activation and recruitment of secretory Rab GTPases to Weibel-Palade bodies. Blood Adv. 5, 5116–5127 (2021).

37. Ishibashi, K., Kanno, E., Itoh, T. & Fukuda, M. Identification and characterization of a novel Tre-2/Bub2/Cdc16 (TBC) protein that possesses Rab3A-GAP activity. Genes Cells 14, 41–52 (2009).

38. Sengeløv, H., Kjeldsen, L. & Borregaard, N. Control of exocytosis in early neutrophil activation. J. Immunol. 150, 1535–1543 (1993).

39. Majumdar, R., Tavakoli Tameh, A., Arya, S. B. & Parent, C. A. Exosomes mediate LTB4 release during neutrophil chemotaxis. PLoS Biol. 19, e3001271 (2021).

40. Arya, S. B., Collie, S. P. & Parent, C. A. The ins-and-outs of exosome biogenesis, secretion, and internalization. Trends Cell Biol. 34, 90–108 (2024).

41. Faurschou, M. & Borregaard, N. Neutrophil granules and secretory vesicles in inflammation. Microbes Infect. 5, 1317–1327 (2003).

42. Cieutat, A. M. et al. Azurophilic granules of human neutrophilic leukocytes are deficient in lysosome-associated membrane proteins but retain the mannose 6-phosphate recognition marker. Blood 91, 1044–1058 (1998).

43. Rada, B. K. et al. Calcium signalling is altered in myeloid cells with a deficiency in NADPH oxidase activity. Clin. Exp. Immunol. 132, 53–60 (2003).

44. Brown, A. C. N. et al. Remodelling of Cortical Actin Where Lytic Granules Dock at Natural Killer Cell Immune Synapses Revealed by Super-Resolution Microscopy. PLOS Biol. 9, e1001152 (2011).

45. Cao, L., Huang, S., Basant, A., Mladenov, M. & Way, M. CK-666 and CK-869 differentially inhibit Arp2/3 iso-complexes. EMBO Rep. 25, 3221–3239 (2024).

46. Hetrick, B., Han, M. S., Helgeson, L. A. & Nolen, B. J. Small molecules CK-666 and CK-869 inhibit actin-related protein 2/3 complex by blocking an activating conformational change. Chem. Biol. 20, 701–712 (2013).

47. Rizvi, S. A. et al. Identification and Characterization of a Small Molecule Inhibitor of Formin-Mediated Actin Assembly. Chem. Biol. 16, 1158–1168 (2009).

48. Serafim, R. A. M. et al. Discovery of a Potent Dual SLK/STK10 Inhibitor Based on a Maleimide Scaffold. J. Med. Chem. 64, 13259–13278 (2021).

49. Collins, S. R. et al. Using light to shape chemical gradients for parallel and automated analysis of chemotaxis. Mol. Syst. Biol. 11, 804 (2015).

50. van den Bos, E., et al. Knockout mouse models reveal the contributions of G protein subunits to complement C5a receptor-mediated chemotaxis. J. Biol. Chem. 295, 7726–7742 (2020).

51. Lämmermann, T. et al. Neutrophil swarms require LTB4 and integrins at sites of cell death in vivo. Nature 498, 371–375 (2013).

52. Kolotila, M. P. & Diamond, R. D. Stimulation of neutrophil actin polymerization and degranulation by opsonized and unopsonized Candida albicans hyphae and zymosan. Infect. Immun. 56, 2016–2022 (1988).

53. Kemp, A. S. & Turner, M. W. The role of opsonins in vacuolar sealing and the ingestion of zymosan by human neutrophils. Immunology 59, 69–74 (1986).

54. Lundgren, S. M. et al. Signaling dynamics distinguish high- and low-priority neutrophil chemoattractant receptors. Sci. Signal. 16, eadd1845 (2023).

55. Zhang, Y. et al. Fast and sensitive GCaMP calcium indicators for imaging neural populations. Nature 615, 884–891 (2023).

56. Rhee, S. G. Regulation of phosphoinositide-specific phospholipase C. Annu. Rev. Biochem. 70, 281–312 (2001).

57. Föger, N. et al. Differential regulation of mast cell degranulation versus cytokine secretion by the actin regulatory proteins Coronin1a and Coronin1b. J. Exp. Med. 208, 1777–1787 (2011).

58. Orci, L., Gabbay, K. H. & Malaisse, W. J. Pancreatic beta-cell web: its possible role in insulin secretion. Science 175, 1128–1130 (1972).

59. Servant, G. et al. Polarization of chemoattractant receptor signaling during neutrophil chemotaxis. Science 287, 1037–1040 (2000).

60. Vargas, A., Roux-Dalvai, F., Droit, A. & Lavoie, J.-P. Neutrophil-Derived Exosomes: A New Mechanism Contributing to Airway Smooth Muscle Remodeling. Am. J. Respir. Cell Mol. Biol. 55, 450–461 (2016).

61. Arya, S. B. et al. Neutrophils secrete exosome-associated DNA to resolve sterile acute inflammation. Nat. Cell Biol. 27, 931–947 (2025).

62. Bailey, T. L. & Gribskov, M. Combining evidence using p-values: application to sequence homology searches. Bioinformatics 14, 48–54 (1998).

