## Extended Data Figures for "Magnetic separation screens identify G alpha 15 as a central regulator of degranulation downstream of neutrophil chemoattractant receptors"

Extended Data Figure 1

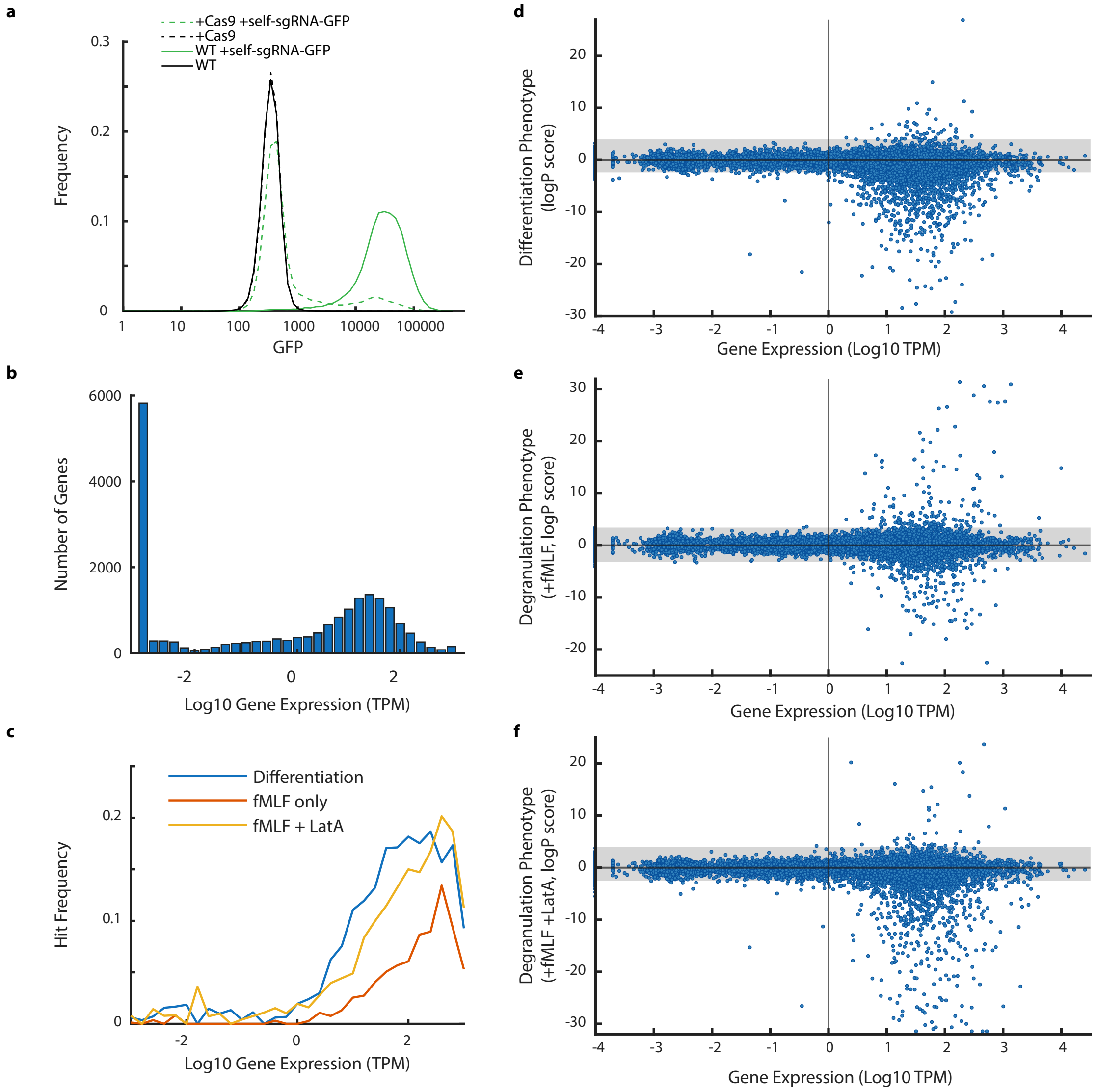


**Extended Data Fig. 1: Validation of Cas9 function and relationship between screen hits and corresponding gene expression levels. a**, GFP expression levels were measured by flow cytometry for dHL-60 cells with stable integration of a Cas9 expression construct and/or a construct expressing GFP and an sgRNA that targets GFP. The reduction of GFP positive cells when both constructs are integrated serves as a measure of Cas9 activity. **b**, Histogram of gene expression levels in log10 transcripts per million (TPM) in dHL-60 cells (data from Rincon et al24). **c**, Frequency of screen hits (defined as passing the 5% FDR threshold) as a function of gene expression level for each of the three genome-wide screens. Gene expression values were binned and the fraction of genes passing the 5% threshold was calculated for each bin. **d-f**, Scatter plots of gene expression level versus the genome-wide screen logP score for the differentiation (**d**), fMLF only degranulation (**e**), and fMLF plus LatA degranulation (**f**) screens. The gray shaded region indicates the logP score range falling below the 5% FDR threshold for each screen. Genes with no detected transcripts are shown at -4 on the gene expression axis.

Extended Data Figure 2

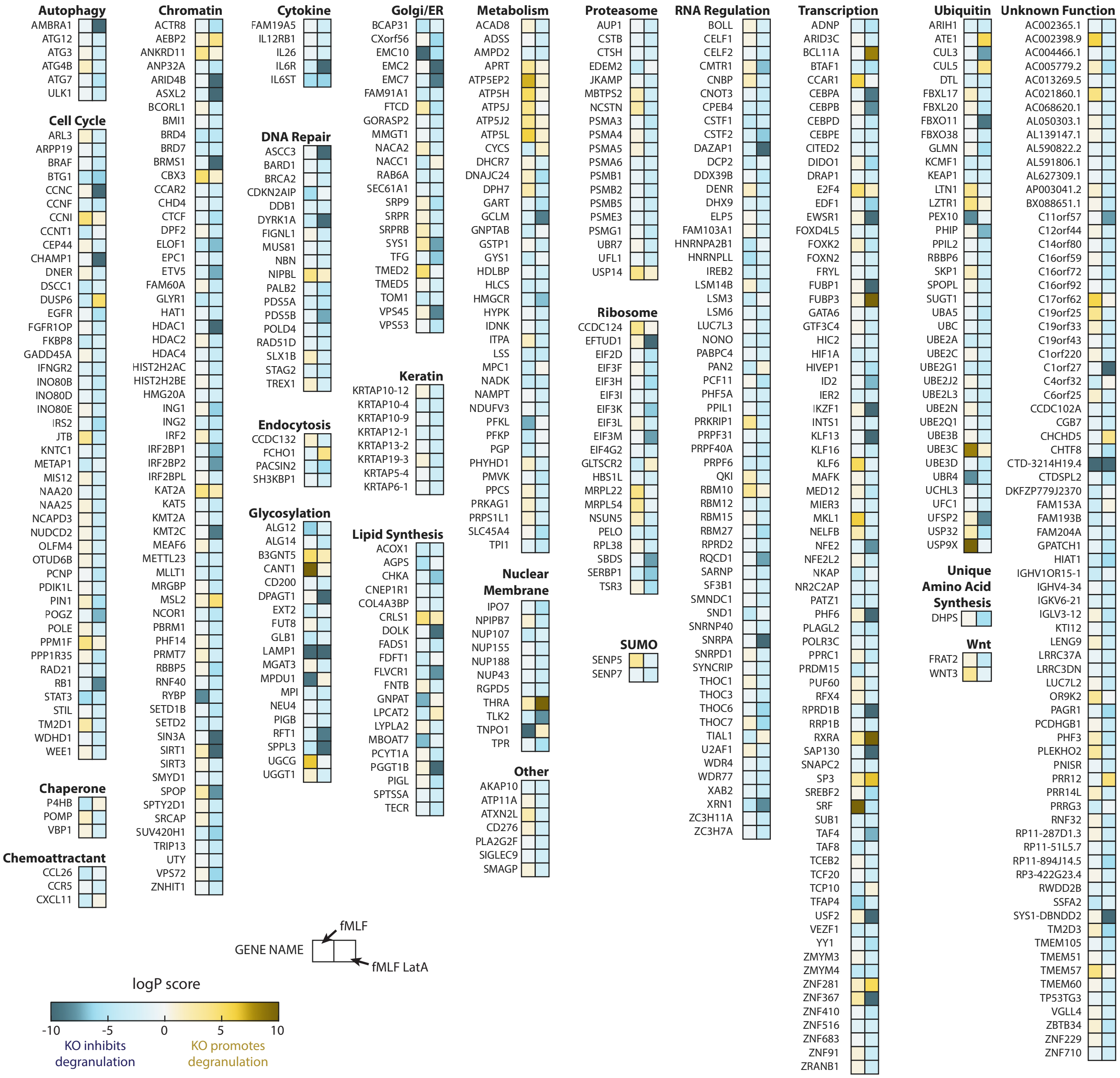


**Extended Data Fig. 2: Continuation of overview of hits from genome-wide degranulation screens**. The schematic is a continuation of the data shown in Fig. 3 and shows logP score heatmaps for genes, organized by functional categories. All genes shown passed a 5% FDR threshold for at least one degranulation screen. Hits for the differentiation screen were excluded. For each gene, the score for the fMLF only screen is shown on the left, and the score for the screen with fMLF and LatA is shown on the right. Functional categories were assigned manually based on prior literature.

### Extended Data Figure 3

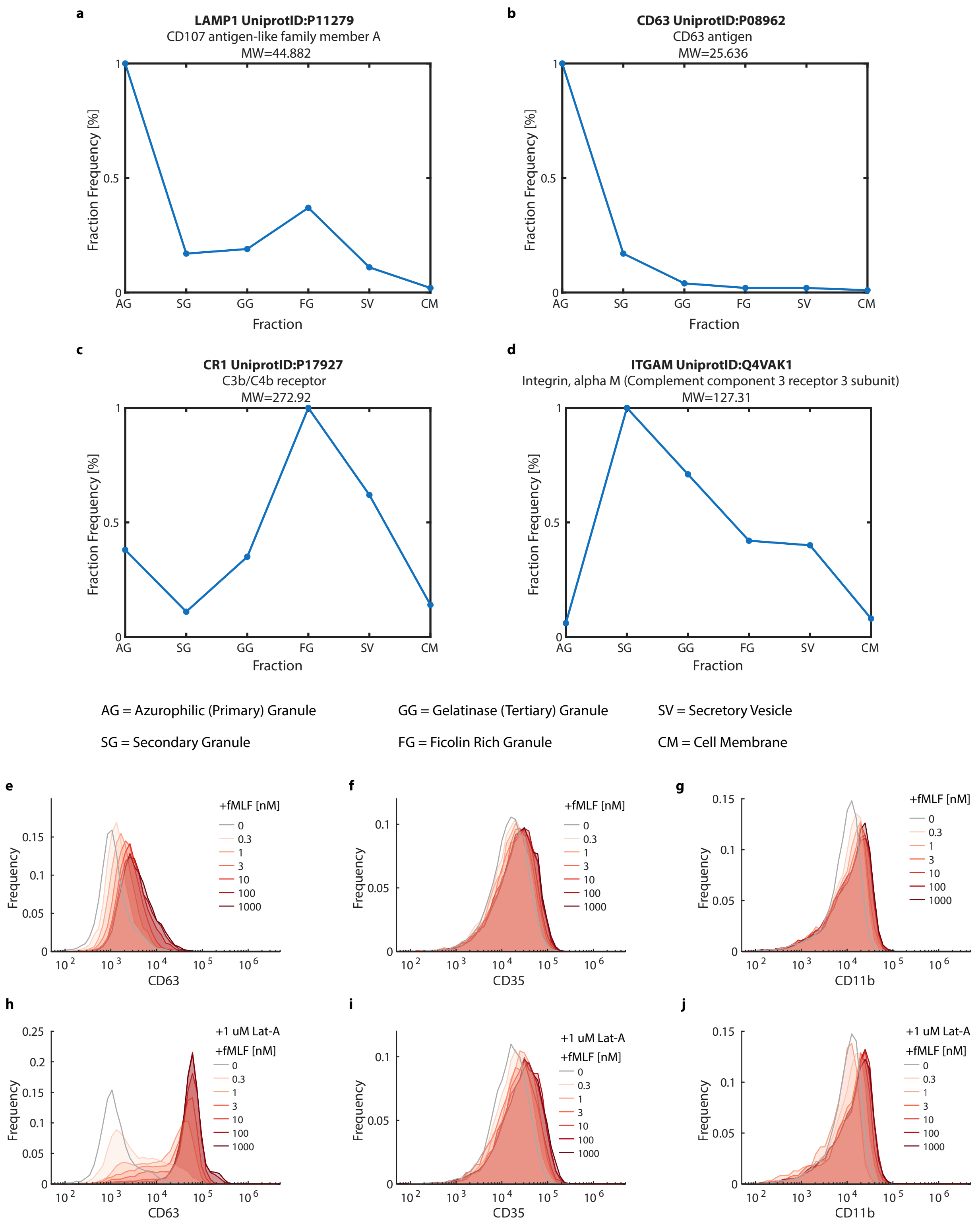


**Extended Data Fig. 3: Granule membrane protein distribution across granule subtypes and measured changes in surface expression in response to fMLF**. **a-d**. Relative abundance of granule marker proteins across granule subtypes. Raw data was obtained from Rørvig et at., 2013[26] based on mass spectrometry measurement of protein abundances in density gradient-separated neutrophil granule fractions. **e-g**. Representative histograms showing cell surface levels of the granule membrane proteins CD63, CD35, and CD11b in dHL-60 cells following stimulation with the indicated concentrations fMLF. **h-j**. Representative histograms showing cell surface levels of the granule membrane proteins CD63, CD35, or CD11b in dHL-60 cells following treatment with the indicated concentrations of fMLF after pretreatment with LatA for 30 minutes.

Extended Data Figure 4

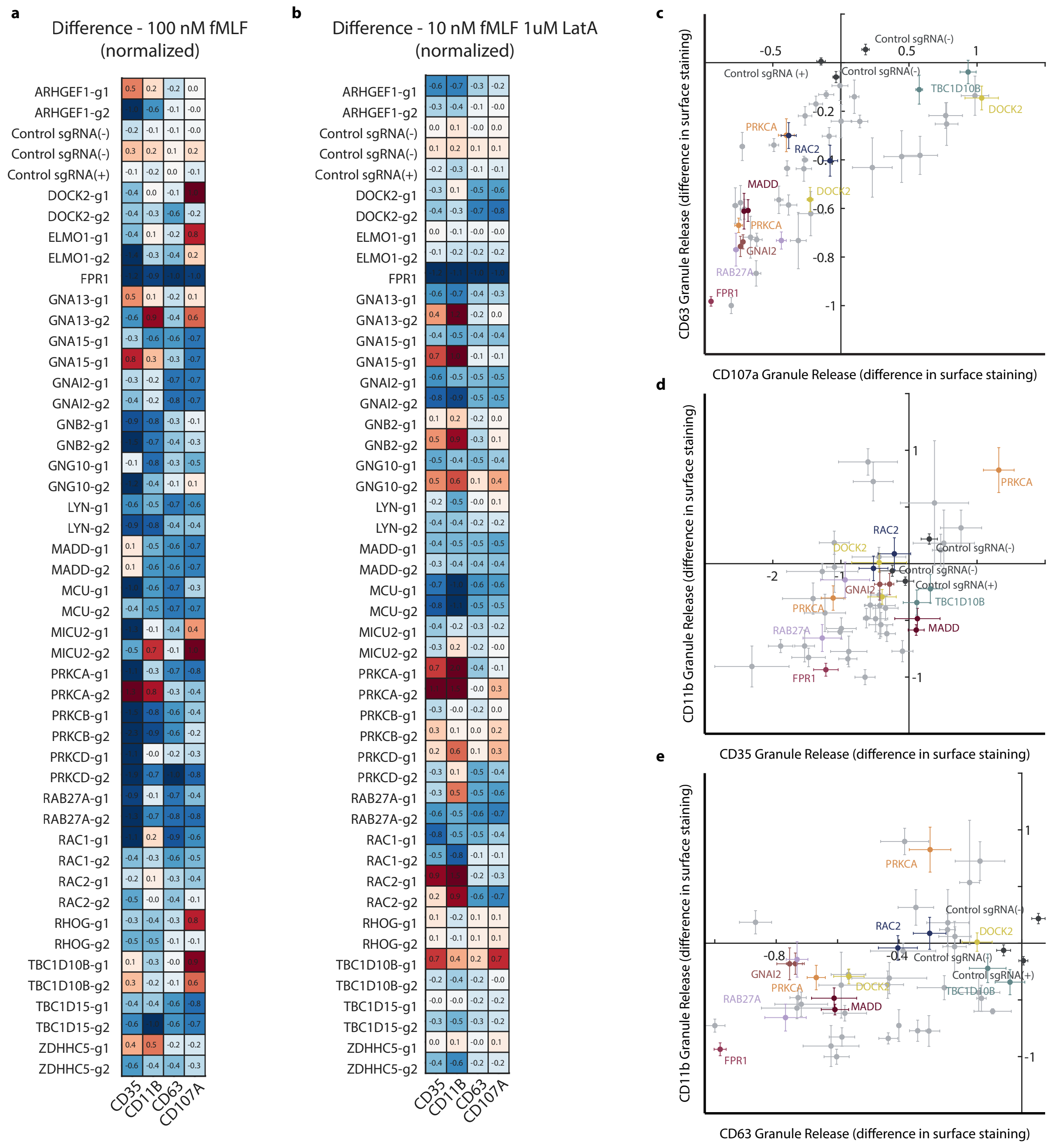


**Extended Data Fig. 4: Analysis of degranulation hits across granule subtypes**. **a, b**. Heatmaps comparing the effects of CRISPRi knockdowns of the selected genes on exocytosis of different granule subtypes in dHL-60s. The effects were measured as the change in cell surface expression of the indicated granule membrane proteins (CD35, CD11b, CD63, and CD107a) after stimulation, then normalized to the mean of the control samples. Cells were stimulated with 100 nM fMLF (**a**) or 10 nM fMLF and 1 uM LatA (**b**). **c-e**, Scatter plots from data in (**a**), comparing the effects of gene knockdowns between granule sub-type exocytosis. Data are normalized to the mean of three control samples. Data points and error bars represent the mean and ±s.e.m. for n=3 independent biological replicates.

Extended Data Figure 5

**a**

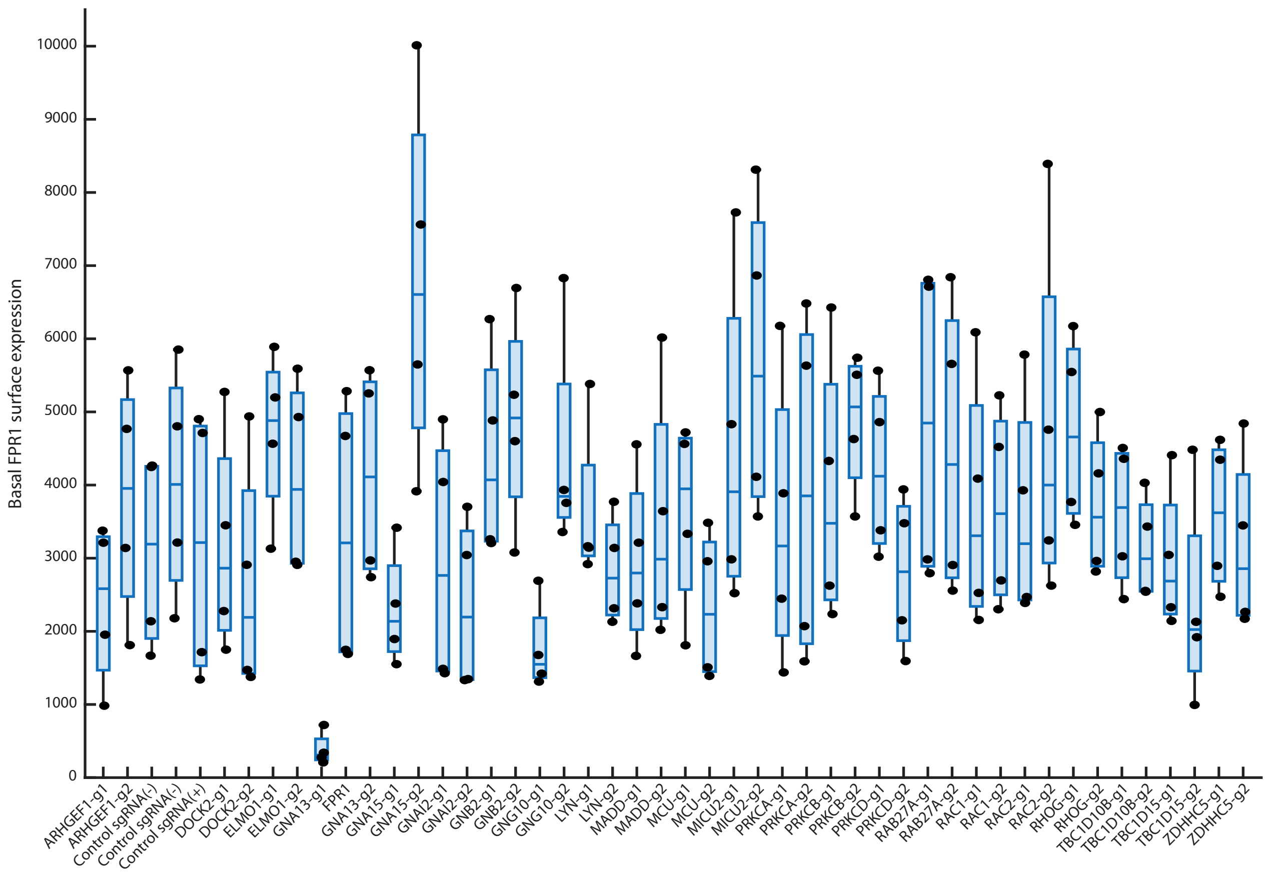


**b**

| | APC-A (FPR1) | PB450-A (CD35) | FITC-A (CD63) | PC7-A (CD107a) | APC-A700-A (CD11b) |
|---|---|---|---|---|---|
| APC-A (FPR1) | 1.000 | 0.000 | 0.000 | 0.000 | 0.000 |
| PB450-A (CD35) | 0.000 | 1.000 | 0.000 | 0.000 | 0.000 |
| FITC-A (CD63) | 0.000 | 0.000 | 1.000 | 0.000 | 0.000 |
| PC7-A (CD107a) | 0.094 | 0.000 | 0.000 | 1.000 | 0.064 |
| APC-A700-A (CD11b) | 0.659 | 0.000 | 0.000 | 0.000 | 1.000 |

**Extended Data Fig. 5: Measurement of cell surface FPR1 protein levels for CRISPRi knockdown cell lines**. **a**, Box plots show cell surface levels of FPR1 measured by flow cytometry (in parallel with measurements of CD35, CD63, CD107a, and CD11b) for the indicated CRISPRi gene knockdown cell lines. For each gene (except FPR1), measurements for two knockdown cell lines are shown, with sg1 on the left and sg2 on the right. These cell lines are the same cell lines used for the experiments in Fig. 4 and Extended Data Fig. 4. **b**, The compensation matrix used to correct cytometry measurements for the experiments in Fig. 4, Extended Data Fig. 4, and panel (**a**).

Extended Data Figure 6

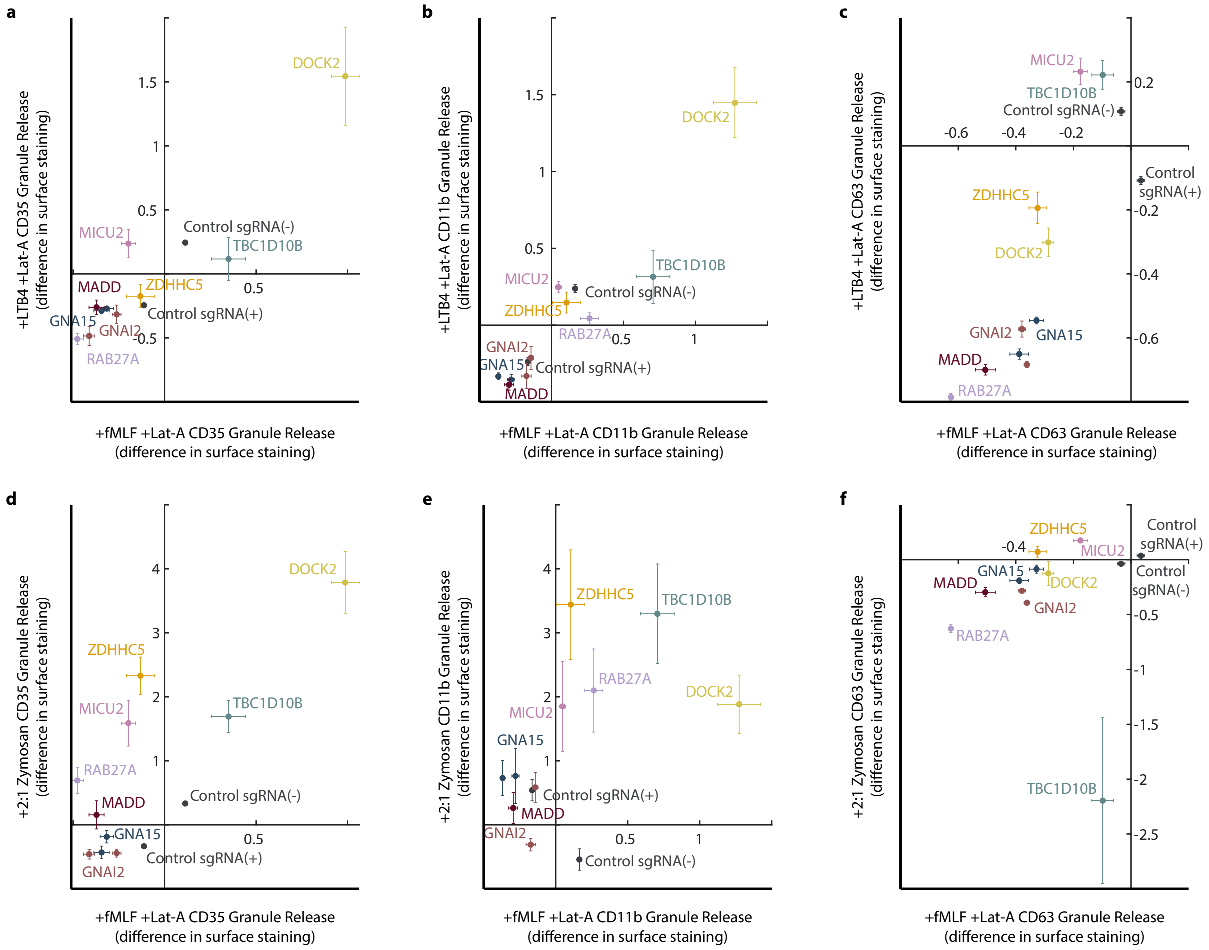


**Extended Data Fig. 6: Comparison of the effects of degranulation hit gene knockdowns for different stimuli**. Scatter plots comparing the effects of gene knockdowns on exocytosis when treated with different stimuli. Each scatter plot shows the data for 10 CRISPRi knockdown cell lines and control cell lines with (+) or without (-) a safe targeting sgRNA. **a-c**, Scatter plots comparing the effects of gene knockdowns on degranulation in response to 100 nM fMLF versus 100 nM LTB4. Degranulation was measured by the change in surface expression of CD35 (**a**), CD11b (**b**), or CD63 (**c**). **d-f**, Scatter plots comparing the effects of gene knockdowns on degranulation in response to 100 nM fMLF versus opsonized Zymosan A (at a ratio of 2 particles per cell) and measured by the change in surface expression of CD35 (**d**), CD11b (**e**), or CD63 (**f**). Data are normalized to the mean of control samples. Data points and error bars represent the mean and ±s.e.m. for n=3 independent biological replicates.

Extended Data Figure 7

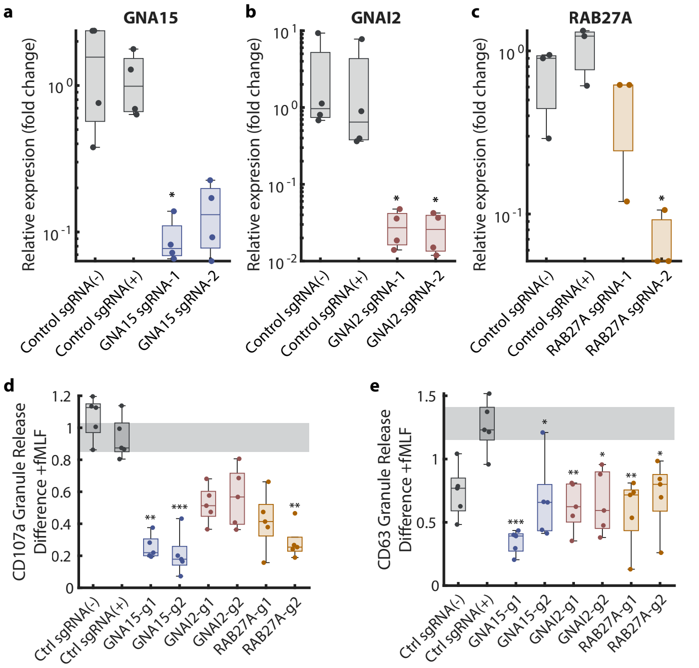


**Extended Data Fig. 7: Assessment of gene knockdown by qPCR and effects of gene knockdowns on degranulation in response to fMLF stimulation (without LatA)**. **a-c**, Quantitative PCR (qPCR) measurement of transcript levels for GNA15 (**a**), GNAI2 (**b**), and RAB27A (**c**) in control or CRISPRi knockdown dHL-60 cell lines. Data were normalized to the mean of the two control cell lines. Dots indicate independent biological replicates. **d,e**, Boxplots quantifying the effect of CRISPRi knockdown of GNA15, GNAI2, or RAB27A on degranulation in response to 100 nM fMLF (without LatA), measured by the difference in cell surface expression of CD107a (**d**) or CD63 (**e**). Data was normalized to the mean of control samples. Dots indicate data from n=5 independent biological replicates. P-values were calculated for each condition against the control sgRNA(+) condition using a nonparametric permutation approach with a Bonferroni correction for multiple comparisons.
